# Loss of SETDB1-mediated H3K9me3 in human neural progenitor cells leads to transcriptional activation of L1 retrotransposons

**DOI:** 10.1101/2025.03.28.645885

**Authors:** Ofelia Karlsson, Ninoslav Pandiloski, Vivien Horvath, Anita Adami, Raquel Garza, Pia A. Johansson, Jenny G Johansson, Christopher H. Douse, Johan Jakobsson

**Affiliations:** Laboratory of Molecular Neurogenetics, Department of Experimental Medical Science and Lund Stem Cell Center, BMC A11, Lund University, 221 84 Lund, Sweden; Laboratory of Epigenetics and Chromatin Dynamics, Department Of Experimental Medical Science and Lund Stem Cell Center, Lund University, Lund, Sweden

## Abstract

Heterochromatin is characterised by an inaccessibility to the transcriptional machinery and is associated with the histone mark H3K9me3. However, studying the functional consequences of heterochromatin loss in human cells has been challenging. Here, we used CRISPRi-mediated silencing of the histone methyltransferase SETDB1 to remove H3K9me3 heterochromatin in human neural progenitor cells. Despite a major loss of H3K9me3 peaks resulting in genome-wide reorganization of heterochromatin domains, silencing of SETDB1 had a limited effect on cell viability. Cells remained proliferative and expressed appropriate marker genes. We found that a key event following the loss of SETDB1-mediated H3K9me3 was the expression of evolutionarily young L1 retrotransposons. Derepression of L1s was associated with a loss of CpG DNA methylation at their promoters, suggesting that deposition of H3K9me3 at the L1 promoter is required to maintain DNA methylation. In conclusion, these results demonstrate that loss of H3K9me3 in human neural somatic cells transcriptionally activates evolutionary young L1 retrotransposons.

## Introduction

At least 50% of the human genome consists of transposable elements (TEs) (1–4). Due to their ability to mobilize, TEs pose a potential threat to genomic integrity and are transcriptionally silenced via epigenetic mechanisms such as histone modifications and DNA methylation (5–10). Long interspersed nuclear elements–1 (L1s) are the most abundant and only autonomously mobilizing family of TEs in humans, accounting for ∼17% of human DNA (1,4,11). Full-length L1s are transcribed from an internal 5′ RNA polymerase II promoter as a bicistronic mRNA encoding two proteins, ORF1p and ORF2p, which are essential for L1 mobilization (12–15). Since L1 elements have colonized the human genome throughout evolution via a copy-paste mechanism in different waves, it is possible to estimate their evolutionary age and assign them to chronologically-ordered subfamilies (16). Hundreds of thousands of L1s are primate specific, and thousands are human specific but only a small fraction of these human-specific full-length L1s are capable of transposing (17,18). Notably, the L1 promoter is bidirectional and L1 antisense transcripts can give rise to chimeric transcripts and act as alternative promoters for protein-coding genes (8,19–21).

L1 transcription is silenced in most human somatic tissues, and transcriptional repression correlates with the presence of heterochromatin marks, including H3K9me3 and CpG DNA methylation (8,9,22,23). In the cells of the human brain, it is well established that CpG DNA methylation plays a critical role in the transcriptional silencing of L1s (6,8,9,23–25). For instance, when DNMT1 is deleted using CRISPR in human neural progenitor cells (NPCs), leading to global DNA demethylation, evolutionarily young full-length L1 elements become transcriptionally active (8). However, it is unclear how DNA methylation is directed to the promoter of evolutionary young L1s in human somatic cells and the interplay between H3K9me3 and DNA methylation in the transcriptional control of L1s is poorly understood.

In this study, we used a somatic human NPC line derived from fetal brain six weeks post-conception (26). This represents a timepoint in post-implantation embryonic development when DNA methylation and other heterochromatin marks have been fully re-established (8,27). This includes the presence of H3K9me3 and DNA methylation over L1s and other TEs (8,23,28), which is established by the KRAB zinc finger (KZNF)-TRIM28 system and other epigenetic TE repressors during early development (29–31). We used CRISPRi to silence SETDB1, a histone methyltransferase associated with these repressors, to study the consequences of H3K9me3 loss at TEs in somatic cells. SETDB1 silencing in hNPCs had a limited effect on cell viability, and the cells remained proliferative and expressed the correct marker genes, despite the widespread loss of H3K9me3 domains. However, SETDB1-mediated loss of H3K9me3 resulted in the transcriptional activation of L1s. Interestingly, this activation correlated with a loss of CpG DNA methylation over the same elements. This finding links the maintenance of DNA methylation at L1s to the presence of H3K9me3. In summary, our results demonstrate the critical role of H3K9me3 in L1 silencing in human neural somatic cells.

## Materials and Methods

### Cell culture

A (male) embryo-derived human neural progenitor cell line (Sai2) was used throughout this manuscript (26). The Sai2 cell line is derived from a small number of cells dissected from fetal brain tissue at Carnegie stage 16, and it has the characteristics of neuroepithelial-like stem cells. The cells were maintained according to the established protocol (32). Cells were grown in poly-L-ornithine (15 ug/ml, Sigma) and laminin 2020 (2ug/ml, Sigma) coated wells. Cells were maintained in culture media DMEM/F12 (Thermo Fisher Scientific) supplemented with N2 (1x, Thermo Fisher Scientific), glutamine (Sigma), Penicillin/Streptomycin (1x, Gibco), B27 (0.05X, Thermo Fisher Scientific), EGF (10ng/ml, Thermo Fisher Scientific) and FGF (10ng/ml, Sigma). Media change was performed daily and cells were passaged every 2-3 days using TrypLE Express (Gibco). 10 *μ*M Y27632 (Rock inhibitor, Miltenyi) was added to the culture media when cells were passaged.

### CUT&RUN

CUT&RUN was performed as previously described by Skene and Henikoff (33). 500.000 hNPCs were washed twice with 20mM HEPES pH 7.5, 150 mM NaCl, 0.5mM spermidine and 1x Roche cOmplete Protease Inhibitor tablet. ConA-coated magnetic beads were activated using 20 mM HEPES pH 7.9, 10 mM, KCl, 1 mM CaCl_2_ and 1 mM MnCl_2_. Cells were attached to preactivated beads and resuspend in 20 mM HEPES pH 7.5, 0.15 M NaCl, 0.5 mM Spermidine, 1x Roche cOmplete protease inhibitors, 0.02% w/v digitonin and 2 mM EDTA together with primary antibody (rabbit anti-H3K4me3, Active Motif 39159; rabbit anti-H3K9me3, Abcam 8898; or goat anti-rabbit IgG, Abcam ab97047) at 1:50 dilution. Bead-bound cells were incubated over night with gentle shake. The following day, the bead bound cells were washed twice with digitonin buffer (20 mM HEPES pH 7.5, 150 mM NaCl, 0.5 mM Spermidine, 1x Roche cOmplete protease inhibitors, 0.02% digitonin) and incubated with pA-MNase diluted in digitonin buffer (700ng/mL) at a rotator for 10 minutes at room temperature. After, cells were washed twice with digitonin buffer, resuspended in digitonin buffer and chilled to 0°C for 5 min. To initiate genomic cleavage, 100mM CaCl_2_ was added, and cells were incubated at 0°C for 30 min. 2x stop buffer (0.35 M NaCl, 20 mM EDTA, 4 mM EGTA, 0.02% digitonin, 50 ng/μL glycogen, 50 ng/μL RNase A) was added to stop the reaction and the cells were incubated at 37°C for 30 min to release genomic fragments. Genomic fragments were purified using PCR clean-up spin column according to manufacturer’s instruction (Macherey-Nagel). Sequencing libraries were prepared using the Hyperprep kit (KAPA) with dual index-adapters (KAPA), pooled and sequenced on a Nextseq500 instrument (Illumina).

### CUT&RUN analysis

Using bowtie2 (--local --very-sensitive-local --no-mixed –no-discordant –phred33 –I –X 700), 2×75bp paired-end CUT&RUN reads were aligned to the human genome (GRCh38) and subsequently converted to BAM files using SAMtools v.1.18 (34). BAM files were then filtered by setting a mapping quality threshold (MAPQ10), which were used to create RPKM (rounds per kilobase million) normalized coverage tracks using bamCoverage (deepTools) v2.5.4 (35). All coverage tracks displayed were visualized in IGV v2.19.1. H3K9me3 peaks were called using SEACR v1.3 (36). Peaks over reference chromosomes longer than 500bp were kept for downstream analyses. Using deepTools’ computeMatrix v 2.5.4 (35), heatmaps matrices were created and visualized using plotHeatmap of the same deepTools package. Finally, we used mergeBed from BedTools v2.31.0 to identify the overlaps of peaks with specific genomic regions, such as evolutionary young L1s or genes. To identify H3K4me3 peaks that overlap with L1 promoters, we used an unbiased k-means strategy to cluster the H3K4me3 signal over all FL L1HS-L1PA3s. This resulted in three clusters. Two of the clusters contained 565 elements with high- or low-confidence H3K4me3 signal. To avoid inflating the number of L1s carrying an H3K4me3 signal, we manually curated the 548 low-confidence elements by inspecting the IGV tracks. This curation process identified 24 elements with a high-confidence H3K4me3 peak at the L1 promoter.

### Lentiviral vector production

Lentivirus was produced as previously described by Zufferey et al (37). Briefly, 293T cells were transfected with third-generation packaging and envelope vectors pMDL (Addgene #12251), psRev (Addgene #12253), and pMD2G (Addgene #12259) with Polyethyleneimine (Polysciences PN 23966) in DPBS (GIBCO). 48h post transfection, lentivirus was harvested by filtering the supernatant and centrifugation at 25,000 × g for 1.5h at 4°C. The virus was resuspended in PBS, aliquoted and stored in −80°C until further use. Lentivirus were in titers of 10^8^-10^9^ as determined by qRT-PCR.

### CRISPRi

CRISPRi constructs were produced as previously described by Johansson et al (38). Single guide sequences were designed to target close to the transcription start site (TSS) using the GPP Portal (Broad institute). The guide sequences were cloned into a lentiviral back bone plasmid containing deadCas9-KRAB-T2A-GFP, a gift from Charles Gersback (Addgene plasmid #71237 RRID:Addgene_71237), using annealed oligos and the BsmBI cloning site. As a control we used one single guide RNA targeting LacZ, a sequence not present in the human genome. Lentivirus was produced for each guide as described above. hNPCs were transduced with MOI 5-7.5 of LacZ, SETDB1, DNMT1, MORC2 and TRIM28 targeting guide RNAs.

For the TRIM28&MORC2 double-CRISPRi we used a double transduction method. Cells were transduced with the previously mentioned deadCas9-KRAB-T2A-GFP lentivirus containing the dCas9 protein, GFP and MORC2 gRNA and simulatniously with lentivirus containing the pLV.U6BsmBI.EFS-NS.H2b-RFPW lentiviral backbone and the gRNA for *TRIM28* and mCherry as a marker. After 10 days expansion, transduced cells were detached using TrypLE Express and resuspended in cell culture media containing Y27632 (10 μM) and propidium iodide (PI, 1:500, BD Bioscience). The GFP^+^ and mCherry^+^ gates were set using untransduced hNPCs and validated via reanalysis of sorted cells (> 95% purity). Transduction efficiency was in the range of 70-95% GFP^+^ cells. Single CRISPRi GFP^+^/PI^-^ and double CRISPRi GFP^+^/mCherry^+^/PI^-^ cells were isolated and pelleted at 400 × g for 10 min. The cell pellets were snap frozen on dry ice and stored in −80°C until further use.

SETDB1 guide 1: AACTCAGGGGTCGGCCTCGA
SETDB1 guide 2: CGAAAGCGAAGGGATAAGGG
DNMT1 guide 1: TGCTGAAGCCTCCGAGATGC
MORC2 guide 1:GCTTCCAAGGACCGGATCGA
TRIM28 guide 1: CCCGCTCGCAGAAAGAGCCG
LACZ guide: TGCGAATACGCCCACGCGAT

### qRT-PCR

Total RNA isolated from hNPCs (RNeasy mini kit, Qiagen) and 250 ng of the isolated RNA was reversed transcribed using the Maxima cDNA synthesis kit (Thermo Fisher Scientific). qRT-PCR was preformed using primers listed below and SYBR green master mix (Roche) on a LightCycler® 480 instrument. Sample were amplified for 40 cycles and cycle values were normalized to ß-actin and HPRT.

Primers:

ß-actin forward: CCTTGCACATGCCGGAG
ß-actin reverse: GCACAGAGCCTCCGCCTT
HPRT forward: ACCCTTTCCAAATCCTCAGC
HPRT reverse: GTTATGGCGACCCGCAG
SETDB1 forward: GACACGTCCAAATATGGGTGC
SETDB1 reverse: ACTGGCTTGAACTGGGTTCC

### Western Blot

For total protein extraction, hNPCs were lysed using RIPA buffer (Sigma-Aldrich) and a complete protease inhibitor cocktail on ice for 30 min, except for the MORC2 protein, where the protein was lysed with 1% (Sigma Aldrich, 71736) instead of RIPA buffer. The lysed cells were pelleted at 17.000 × g for 20 min at 4°C. For total histone protein extraction, we used a Histone Extraction kit (Abcam) according to manufacturer’s instructions. 2,5 ug of extracted total protein or 2,5 ug of extracted histone proteins were mixed with Novex LDS 4x loading dye (Thermo) and reducing agent (Thermo) before being boiled at 95°C for 5 minutes. Samples were loaded onto a 4-12% SDS/PAGE gel and run at 150V for 0.5h. The proteins were transferred to a PVDF membrane using a Transblot-Turbo transfer system (BioRad). The membrane was washed two times in TBS with 0.1% Tween (TBST) before being blocked in TBST with 5% skimmed milk for 1h. The membrane was incubated overnight at 4°C with primary antibody mouse anti-SETDB1 (Thermo Fisher MA5-15722, 1:2000), rabbit anti-H3K9me3 (Abcam 8898, 1:20.000) rabbit anti-histone 3 (Abcam 18521, 1:250.000), rabbit anti-MORC2 (A300-149A, Bethyl; 1:1000 dilution), rabbit anti-TRIM28 (Abcam, ab10484; 1:1000 dilution) diluted in TBST with 5% skimmed milk. The following day, the membrane was washed three times with TBST before being incubated with secondary antibody HRP-conjugated anti-mouse antibody (Santa Cruz Biotechnology, 1:5.000), HRP-conjugated anti-rabbit antibody (Santa Cruz Biotechnology, 1:10.000 or 1:50.000) or HRP conjugated anti-ß-actin (Sigma, A3854, 1:50.000) diluted in TBST with 5% skimmed milk for 2h in room temperature. The membrane was washed three times with TBST and once with TBS before protein was detected by chemiluminescence using ECL Prime Western blotting detection reagent (Cytiva). The signal was detected using a ChemiDoc system (BioRad).

### Immunocytochemistry: SOX2 and NESTIN

hNPCs were washed three times with DPBS and fixed using 4% paraformaldehyde for 15 min at room temperature, followed by three more washed with DPBS. After incubation in blocking solution (KPBS + 0.25% Triton X-100 with 5% normal donkey serum) for 1h at room temperature, the cells were incubated with primary antibodies SOX2 (R&D Systems, AF2018, 1:100) and NESTIN (Abcam, AB176571, 1:100) diluted in blocking solution overnight at 4°C. After being washed twice with TKPBS and once with TKPBS + 5% NDS, cells were incubated with secondary antibodies Alexa647-donkey anti-rabbit (Jackson Immuno Research 711-605-152, 1:500) and Cy3-donkey anti-goat (Jackson Immuno Research 705-165-533 003, 1:500) diluted in blocking solution for 2h at room temperature following incubation with DAPI (Sigma, 1:1000) for 5 min. The cells were rinsed two times with KPBS and imaged using a Leica microscope (model DMI6000 B). Images were cropped and adjusted using ImageJ Fiji.

### Immunocytochemistry: H3K9me3

hNPCs plated in poly-L-ornithine (15 ug/ml, Sigma) and laminin 2020 (2ug/ml, Sigma) coated 24 well ibidi plates were rinsed three times with DPBS before being fixed with 4% paraformaldehyde at room temperature for 15 min. The cells were washed once with KPBS before being blocked in TKPBS with 5% NDS for 1 h at room temperature. After, the cells were incubated with primary antibody rabbit anti-H3K9me3 (Abcam 8898, 1:200) diluted in TKPBS + 5% NDS overnight at 4°C. The following day, cells were washed three times with TKPBS before being incubated with secondary antibody Cy5-donkey anti-rabbit (Jackson Immuno Research,1:500) diluted in TKPBS + 5% NDS at room temperature for 1h. The cells were washed three times with KPBS and incubated with DAPI (1:1000) diluted in TKPBS at room temperature for 5 min. The cells were washed twice with KPBS before being imaged using confocal Microscope Leica TSC SP8. Images were cropped and adjusted using ImageJ Fiji.

### EdU proliferation assay

To study cell proliferation we used the Click IT^TM^ Plus EdU Alexa Fluor^TM^ 647 Imaging Kit (Invitrogen, C10640). GFP^+^ SETDB-CRISPRi and Control-CRISPRi cells were FACS sorted 10 days after transduction and re-plated for the EdU assay. After 24h incubation with EdU (10 µM), cells were fixed in 4% paraformaldehyde. After fixing, cells were stained for Alexa647 and Hoechst following the manufacturer’s instructions. Cells were imaged and analyzed using an Operetta CSL high-content analysis system.

### Bulk RNA sequencing

Total RNA was isolated from hNPCs using the RNEasy mini kit (Qiagen) with on column DNAse treatment. Libraries for bulk RNA sequencing were generated using Illumina TruSeq Stranded mRNA library prep kit (with poly-A selection) optimized for long fragments. Sequencing was performed on a NovaSeq6000 (paired end 2 × 150bp).

### Bulk RNA sequencing analysis

Paired-end 150bp RNAseq reads were mapped to the reference human genome (GRCh38) using STAR aligner verson 2.6.0 (39) with gencode v38 as a guide (--sjdbGTFfile). Unique mapping (described below) was performed to investigate the expression of individual transposable elements (TEs). Multi mapping (described below) was performed to investigate the expression of genes and TEs at a subfamily level.

#### Unique mapping

Reads were allowed to align to a single locus (--outFilterMultimapNmax 1) with a mismatch ratio up to 0.03 (–outFilterMismatchNoverLmax 0.03). Sorted alignment files as output by STAR were indexed using SAMtools v1. 18 (34). Normalized coverage tracks (bigwig files) using RPKM were created using deeptools bamCoverage v2.5.4 (35). FeatureCounts v1.6.3 (40) was used to quantify the expression of individual TEs using a curated GTF file provided by the creators of TEtranscripts. FeatureCounts was ran on default parameters using –s2 to force the strandedness of the reads to the feature.

#### Differential expression analyses

DESeq2 v1.38.3 (41) with default parameters was used to perform differential expression analyses for all bulk RNAseq datasets using the read count matrix obtained with featureCounts. Mean plots were created using median-of-ratio normalized counts per condition, adding a pseudo count of 0.5, and log2 transformed values. All pvalues and p adjusted values calculated by DESeq2 are two-sided.

To obtain distribution of the expected upregulated TEs, the RepeatMasker annotation was sampled with the exact number of the observed upregulated TEs, without replacement. The number of sampled TE was then compared to the observed values and scored accordingly (+1 if expected > observed; +0 if expected ≤ observed). This then was repeated 50.000 times to calculate a double-sided p-value.

To quantify and normalise the strand-specific expression of L1s, the reads were normalised using gene-derived medians to calculate the median ratio. p-values were calculated using the Mann–Whitney U-test, a non-parametric statistical test used to compare the non-normal distributions of two independent groups. When calculating the p-value for the differences between conditions, each L1 element was considered an independent replicate. In the analysis we used the FL-L1s that were upregulated and found in at least one of the two experimental conditions (two different gRNAs). This resulted in 241 FL-L1s being included in the analysis. Gene-ontology overrepresentation test of biological processes (enrichGO) was performed using clusterProfiler v4.8.2 over differentially expressed genes (|LFC| > 1, two-sided padj < 0.05). To measure sense and antisense transcription over evolutionary young L1s, featureCounts was used specifying the -s2 and -s1 parameter, respectively. Gene expression size factors were further used to normalize the reads by median-of-ratios normalization approach.

### Oxford Nanopore DNA sequencing

High molecular weight DNA was extracted from hNPCs using the Nanobind HMW DNA extraction kit (PacBio) according to the manufacturer’s instructions. DNA quality and concentration was measured using Nanodrop and the dsDNA HS assay on a Qubit (Thermo Fisher Scientific). Genomic fragments <5kb were eliminated using the Short Read Eliminator XS kit (PacBio) and remaining genomic fragments were sheered to 10kb using a g-tube kit (Covaris) following manufacturer’s instructions. Libraries were prepared using the SKQ-LSK114 kit (Oxford Nanopore Technologies) with 1 µg DNA input and loaded onto R10.4.1 flow cells. Sequencing was performed on an Oxford Nanopore Technologies PromethION 2 solo for 72h following manufacturers guidelines. To obtain the data, dorado (v4.3.0)(RRID: <u>SCR_025883</u>; https://github.com/nanoporetech/dorado) modified basecalling was performed over the pod5 files, of the 5mCG_5hmCG super accurate basecalling model. Basecalled files were mapped to the human genome (GRCh38) using minimap2 v2.25 (42). Alignment files (BAM) sorted and indexed using SAMtools, and methylation matrices were created using the segmeth tool from MethylArtist v1.2.6 (43). Methylartist segplot and locus tools were further employed for all downstream visualization, either using methylation matrices or BAM files as input. To statistically compare the difference in DNA methylation between control and CRISPRi conditions, Wilcoxon signed-rank test was used. Methylartist composite plots were created using L1HS and L1PA2 consensus sequences to plot FL L1HS and FL L1PA2-L1PA3 elements, respectively. By default the tool uses a maximum of 200 elements per plot, therefore if elements of interest surpassed that threshold, 200 of them were randomly selected with “shuf-n 200”.

Since the modified basecalling model also provides information about 5hmC methylation, the same BAM files were used to asses the changes in 5hmC methylation over promoter regions of L1s. To visualize the data and statistically compare the differences in 5hmC methylation, identical approaches were used as above with the only exception of specifying the “-m h” flag to obtain 5hmC methylation over the chosen regions. The y-axes of the locus plots were altered using “-ymin 0 -ymax 0.2”.

## Results

### L1s are covered by H3K9me3 in hNPCs

To investigate the distribution of H3K9me3 heterochromatin in hNPCs we performed CUT&RUN epigenomic analysis. We used a hNPC line that shares characteristics with neuroepithelial-like stem cells **(Fig. 1A)**. This cell line was derived from fetal brain at six weeks post-conception (26), a time point in post-implantation embryonic development when heterochromatin and DNA methylation patterns have been fully re-established.

**Figure 1.**
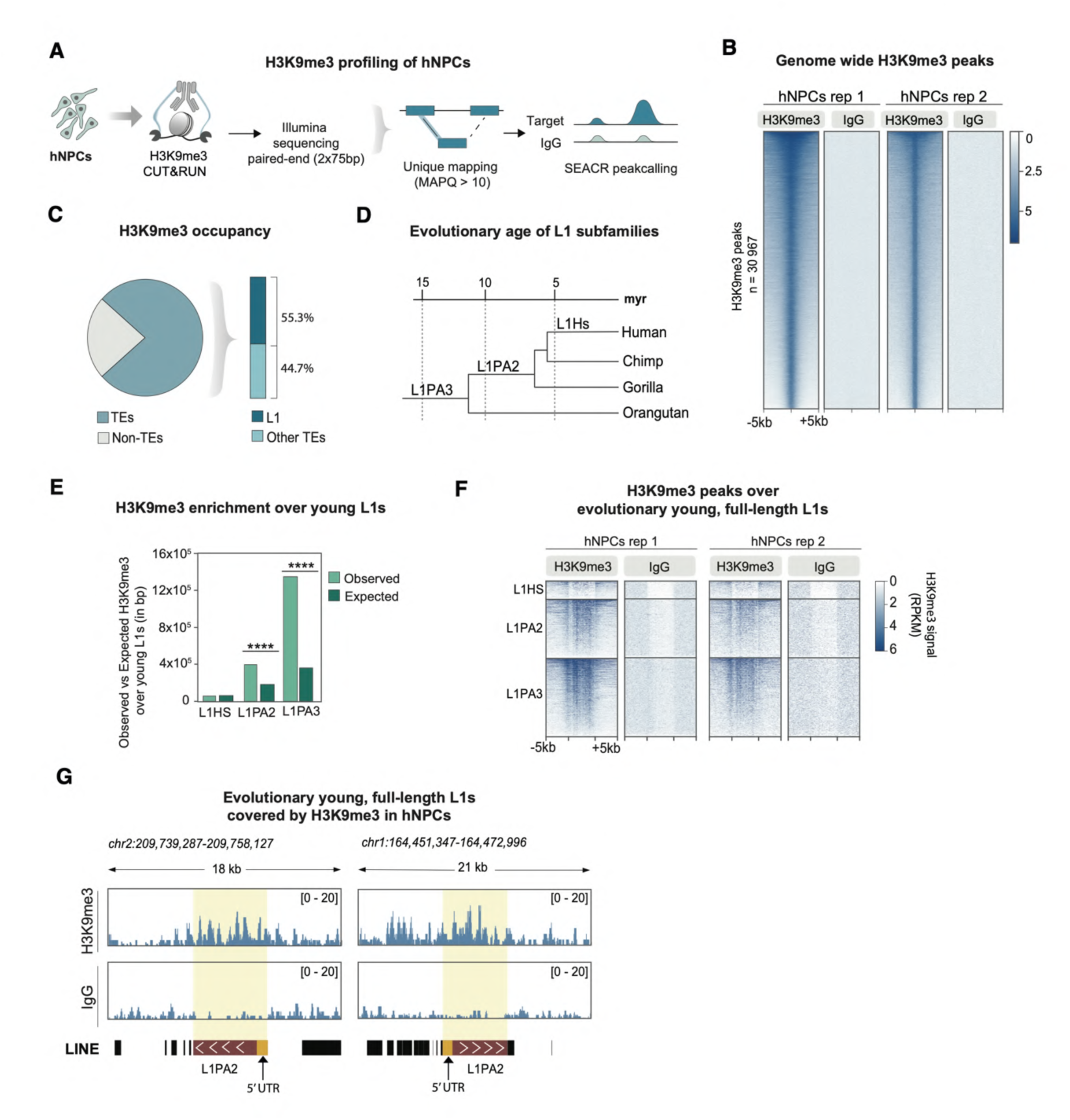
Evolutionary young, full-length L1s are covered by H3K9me3 in hNPCs. **A** Schematic illustrating CUT&RUN epigenomic profiling of H3K9me3 in hNPCs. **B** RPKM normalized heat maps showing genome-wide CUT&RUN signal enrichment of H3K9me3 and non-targeting control IgG in hNPCs (n=2). **C** Plots showing the genome-wide distribution of H3K9me3 CUT&RUN peaks in hNPCs, categorized by TEs, non-TE (pie chart); L1 elements and other TEs (bar chart). **D** Phylogenetic tree of evolutionarily young L1 subfamilies. **E** Barplot showing H3K9me3 enrichment over evolutionary young L1s **F** RPKM normalized heat maps showing CUT&RUN signal of H3K9me3 in hNPCs and their respective IgG controls, over full-length L1HS to L1PA3 elements sorted by evolutionary age (n=2). **G** Genome Browser tracks showing RPKM normalized H3K9me3 CUT&RUN data over two full-length L1PA2 elements in hNPCs.

Due to the high degree of sequence similarity among L1s, short read sequencing data results in a large proportion of ambiguously mapping reads. To avoid signal amplification due to multimapping artifacts over these elements, we used a strict unique mapping approach to examine individual L1 loci (**Fig. 1A**). With this bioinformatic approach, it is possible to analyze the epigenetic status of most L1s, except for some of the evolutionarily-youngest elements, where we rely on the flanking regions **(Sup Fig 1A)**. This approach allows us to discriminate reads without ambiguity, and trace the epigenetic modification to a unique locus in the human genome.

We found that H3K9me3 is highly abundant in hNPCs and peak calling identified 30.967 regions in the human genome covered by H3K9me3 **(Fig. 1B, Sup Fig. 1B)**. More than three quarters of the peaks (76.9%) could be confidently mapped to TEs, and among these, L1s represented the most abundant TE class (55.3%) **(Fig. 1C).** We found that H3K9me3 was particularily enriched over evolutionary young L1s (**Fig. 1D-E**). Almost all full-length (>6kb) evolutionarily-young elements, including human-specific (L1HS) and hominoid-specific (L1PA2 and L1PA3) copies, were covered by H3K9me3 in hNPCs **(Fig. 1F-G)**. Notably, the majority of these full-length L1 elements contain an intact 5’ promoter and can be transcribed, and some even retain the capacity to retrotranspose (11,18). These data demonstrates that hNPCs represent a useful model for investigating epigenetic mechanisms of L1 regulation, including the presence of heterochromatin, in human somatic neural cells.

### CRISPRi-based silencing of SETDB1 in hNPCs results in a global reorganization of H3K9me3

The deposition of H3K9me3 at TEs has been linked to the histone methyltransferase SETDB1 (also known as ESET) (44–46), which is highly expressed in hNPCs and the developing and adult human brain **(Sup Fig. 2A-C)**. To study the effects of the loss of SETDB1-mediated H3K9me3, we developed a CRISPRi-based approach to transcriptionally silence SETDB1 in hNPCs (SETDB1-CRISPRi) **(Fig. 2A)**. Two guide RNAs were designed to target near the transcription start site (TSS) of the SETDB1 gene. As a control, we used a guide RNA targeting LacZ, a sequence not found in the human genome. SETDB1 and control guide RNAs were separately cloned into lentiviral vectors expressing a catalytically inactive Cas9 (dCas9) fused to a Krüppel-associated box (KRAB) transcriptional repressor domain (38). Transduced cells were expanded for 10 days prior to FACS isolation of GFP^+^ cells carrying the CRISPRi construct. Lentiviral transduction of hNPCs using the SETDB1-CRISPRi vector resulted in efficient, near-complete silencing of SETDB1 at transcript and protein level using both gRNAs, as determined by qRT-PCR, Western blot and RNA sequencing (**Fig. 2B-D, Sup Fig. 3A-C)**. Since CRISPRi works by transcriptionally silencing its target TSS through heterochromatin remodeling via KRAB repressor binding, it was surprising that SETDB1 silencing could be achieved, since the KRAB system depends on this enzyme for H3K9me3 deposition (47). To investigate the mechanism behind this, we performed CUT&RUN profiling of H3K4me3 and H3K9me3 of SETDB1-CRISPRi hNPCs. Analysis of the active promoter mark H3K4me3 revealed a complete loss of H3K4me3 over the SETDB1 TSS following SETDB1-CRISPRi, confirming the loss of transcriptional activity at the SETDB1 promoter. However, this was not accompanied by an accumulation of the repressive histone mark H3K9me3 over the SETDB1 TSS (**Fig. 2D)**. These results show that the CRISPRi machinery works efficiently without H3K9me3 deposition, at least at this locus.

**Figure 2.**
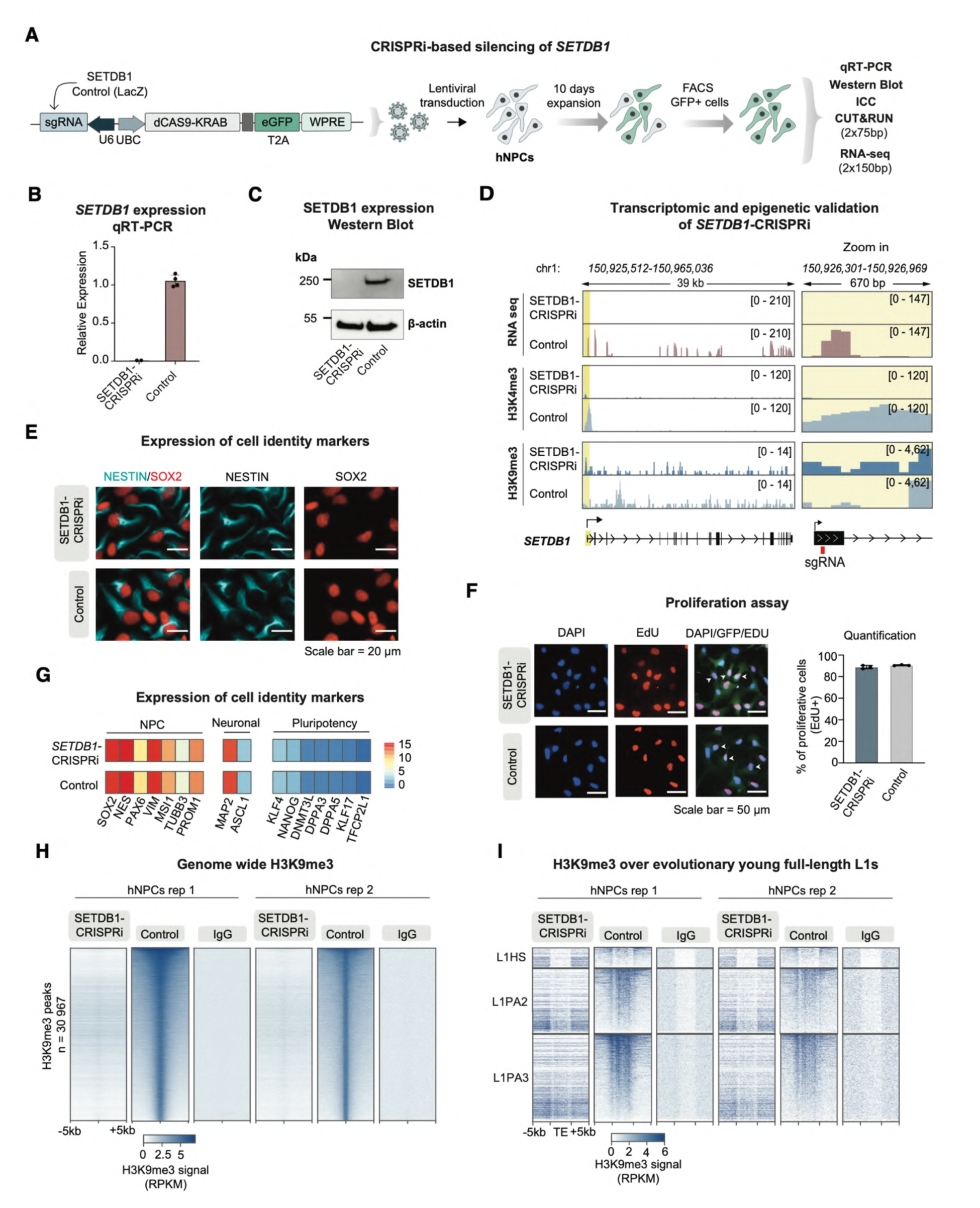
SETDB1-CRISPRi results in loss of H3K9me3 in hNPCs. **A** Schematic of the CRISPRi construct and workflow for CRISPRi-based silencing of SETDB1 in hNPCs. **B** qRT-PCR analysis of SETDB1 expression after CRISPRi silencing in hNPCs. Results are shown as means with standard deviation (n=2 and n=4 respectively). **C** Western blot analysis of SETDB1 protein levels in SETDB1-CRISPRi and control hNPCs relative to β-actin protein levels. **D** RPKM normalized genome browser tracks showing: expression of SETDB1 in SETDB1-CRISPRi and control hNPCs (top). Epigenetic changes as a result of the CRISPRi approach as determined by CUT&RUN profiling of H3K4me3 (middle) and H3K9me3 (bottom). **E** Immunocytochemistry of NESTIN (cyan) and SOX2 (red) in SETDB1-CRISPRi and control hNPCs. Scale bar = 20 µm. **F** EdU proliferation assay showing cell proliferation in SETDB1-CRISPRi and Control hNPCs. Scale bar=50 µm. **G** Heat map showing mean expression of NPC, neuronal, and pluripotency gene markers in SETDB1-CRISPRi (n=2) and control (n=4) hNPCs as determined by bulk RNA sequencing. **H** Heat maps illustrating genome-wide RPKM normalized CUT&RUN signal of H3K9me3 and non-targeting control IgG in SETDB1-CRISPRi (n=2) and control (n=2) hNPCs. **I** Heat maps illustrating RPKM normalized CUT&RUN signal of H3K9me3 and non-targeting control IgG in SETDB1-CRISPRi (n=2) and control (n=2) hNPCs over full-length L1HS-L1PA3 elements.

**Figure 3.**
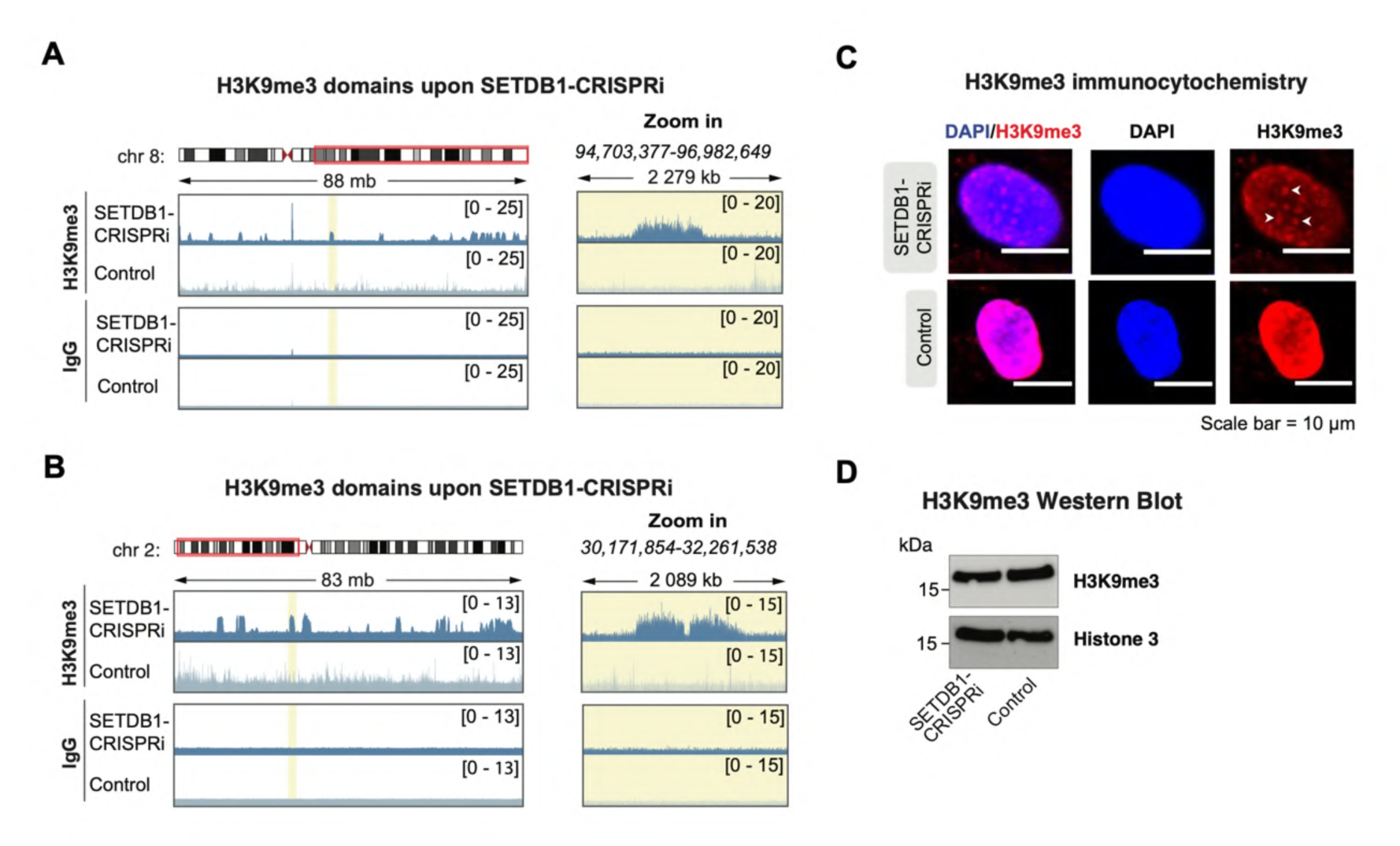
SETDB1-CRISPRi results in reorganisation of H3K9me3 in hNPCs. **A** Genome browser track of chromosome 8 showing RPKM normalized CUT&RUN signal of H3K9me3 in SETDB1-CRISPRi and control hNPCs. Right panels show zoom in of H3K9me3 dense areas highlighted in yellow. **B** Genome browser track of chromosome 2 showing RPKM normalized CUT&RUN signal of H3K9me3 in SETDB1-CRISPRi and control hNPCs. Right panels show zoom in of H3K9me3 dense areas highlighted in yellow. **C** Immunocytochemistry of H3K9me3 (red) and nuclear marker DAPI (blue) in SETDB1-CRISPRi and control hNPCs. H3K9me3 foci are indicated with white arrows. Scale bar = 10 µm. **D** Western blot analysis of H3K9me3 protein levels in SETDB1-CRISPRi and control hNPCs relative to Histone 3 protein levels.

The SETDB1-CRISPRi hNPCs remained viable despite the absence of SETDB1 expression and no changes in morphology were observed compared to control cells **(Fig. 2E, Sup Fig 3D)**. Previous studies in mouse model systems have described H3K9me3 as important for maintaining the cellular identity of neural cells and silencing lineage-inappropriate genes (48). However, in SETDB1-CRISPRi hNPCs proliferation remained unchanged, as monitered by EdU incorporation assays **(Fig. 2F, Sup Fig3E)**. Expression of NPC lineage markers did not differ from that of control hNPCs, as determined by bulk RNA sequencing and neuronal and pluripotency markers were not activated, at least over the 10-day timecourse of the experiment (**Fig. 2G, Sup Fig. 3F**). Furthermore, immunofluorescence staining for the NPC markers SOX2 and NESTIN showed no obvious differences in SETDB1-CRISPRi hNPCs compared to controls (**Fig. 2E, Sup Fig. 3D**). To further investigate how silencing of SETDB1 affects the cellular identity of hNPCs, we performed differential gene expression analysis. We found that 748 genes were upregulated and 1202 genes were downregulated in SETDB1-CRISPRi hNPCs compared to controls (|LFC| > 1, two-sided padj < 0.05, DESeq2) **(Sup Fig. 3G)**. Gene-ontology overrepresentation test of differentially expressed genes (|LFC| > 1) in SETDB1-CRISPRi hNPCs revealed an enrichment of terms related to cell adhesion and synapse assembly among the upregulated genes, and regulation of actin filament organization among the downregulated genes **(Sup Fig. 3H).** Taken together, these results demonstrate that CRISPRi-mediated silencing of SETDB1 results in modest transcriptional changes, but that these do not appear to have a major impact on the cellular identity of hNPCs.

To investigate the role of SETDB1 in depositing H3K9me3 in human NPCs, we performed CUT&RUN profiling of H3K9me3 in SETDB1-CRISPRi and control cells. This revealed a genome-wide loss of the majority of H3K9me3 peaks. Of the 30.967 H3K9me3 peaks found in control hNPCs, 85.6 % were lost in SETDB1-CRISPRi hNPCs **(Fig. 2H).** This included a profound loss of H3K9me3 over evolutionarily-young full-length L1 elements **(Fig. 2I)**. However, not all H3K9me3 was lost. Surprisingly, we found the appearance of large ectopic regions of H3K9me3, mainly at centromeres and telomeres, but also scattered throughout the genome including regions flanking some of the FL-L1s **(Fig. 3A,B, Sup Fig. 4A)**. The appearance of these ectopic regions of H3K9me3 was also visible when performing immunocytochemistry for H3K9me3 in SETDB1-CRISPRi hNPCs **(Fig. 3C, Sup Fig. 4B).** We found no changes in global H3K9me3 levels using WB analysis **(Fig. 3D, Sup Fig. 3C).** The underlying mechanism for the appearance of these broad ectopic H3K9me3 domains remains unclear but is likely a result of the activity of other H3K9me3 methyltransferases (49) and the complete heterochromatin reorganization that occurs in SETDB1-CRISPRi hNPCs.

**Figure 4.**
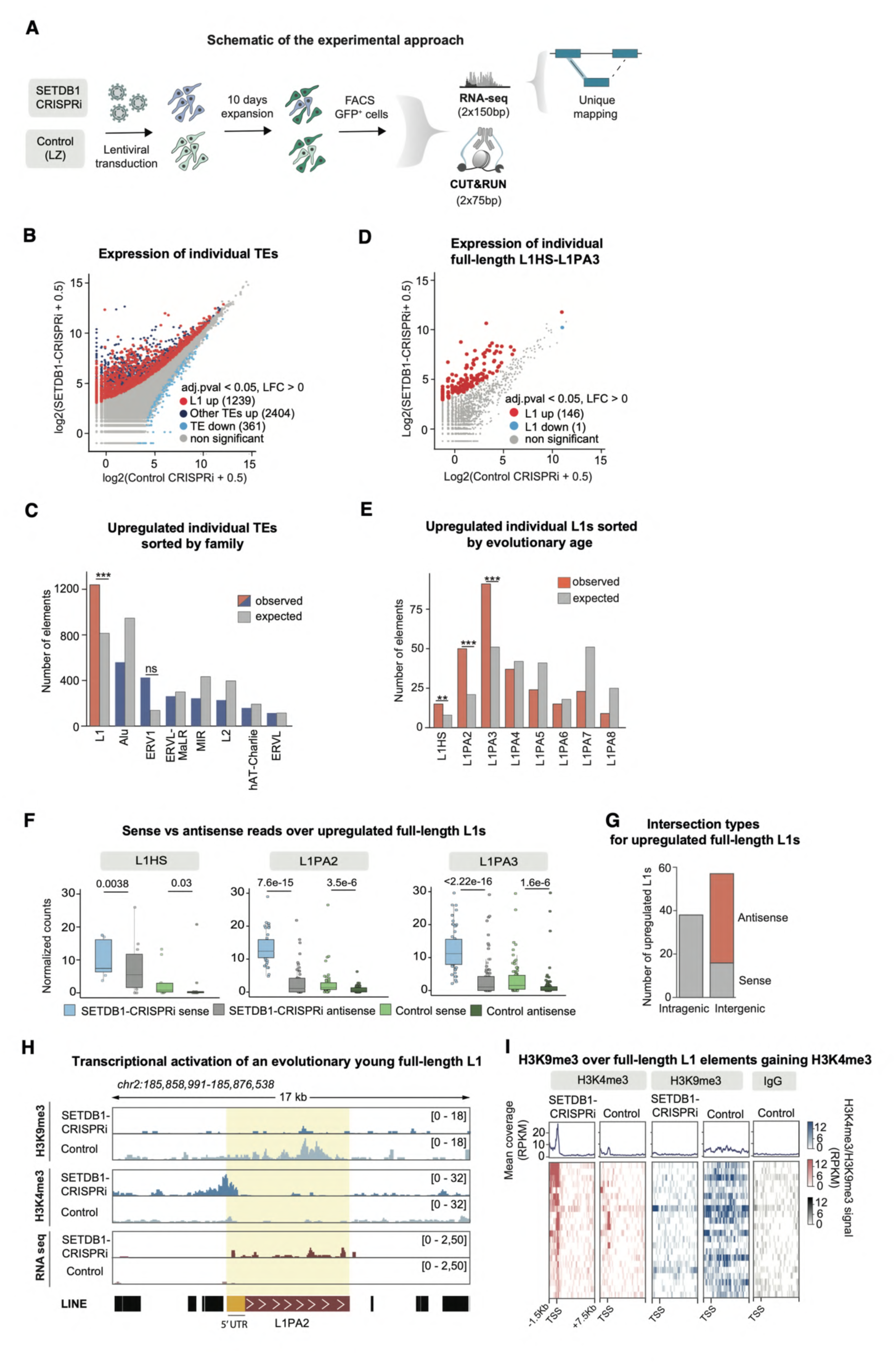
Evolutionarily-young, full-length L1s are transcriptionally activated upon loss of H3K9me3 in hNPCs. **A** Workflow for CUT&RUN and RNA sequencing analysis of SETDB1-CRISPRi and control hNPCs. **B** Mean plot of uniquely mapped bulk RNA sequencing reads for SETDB1-CRISPRi hNPCs (n=2) compared to control (n=2) over individual TEs categorized into L1 elements and other TEs. Log_2_ fold change (LFC) >0, two-sided padj 0.05 calculated with DESeq2. **C** Observed vs expected number of individual, transcriptionally activated elements per TE family in SETDB1-CRISPRi hNPCs (n= 2) compared to control (n= 2). Pvalues were calculated using bootstrap-based empirical two-sided pvalue calculation. **D** Mean plot of uniquely mapped bulk RNA sequencing reads for SETDB1-CRISPRi hNPCs (n=4, pooled data from both gRNAs) compared to controls (n=4) over individual full-length (>6 kb) L1HS-L1PA3 elements. LFC > 0, two-sided padj < 0.05 calculated with DESeq2. **E** Observed vs expected number of transcriptionally activated L1 elements sorted by evolutionary age in SETDB1-CRISPRi (n=2) compared to control hNPCs (n=2). Pvalues were calculated using bootstrap-based empirical two-sided pvalue calculation. **F** Normalized read counts in sense and antisense for upregulated, full-length L1HS-L1PA3 elements in SETDB1-CRISPRi (n=2) vs control hNPCs (n=4). Statistical significance was calculated with Mann-Whitney U test. **G** Number, genomic location and direction of upregulated full-length (>6 kb) L1HS-L1PA3 elements in SETDB1-CRISPRi (n=2) compared to control hNPC (n=2) **H** RPKM normalized genome browser tracks showing: CUT&RUN analysis of H3K9me3 (top), H3K4me3 (middle) and bulk RNA sequencing (bottom) in SETDB1-CRISPRi and control hNPCs. **I** RPKM normalized heat maps showing CUT&RUN signal of H3K9me3 over full-length (>6 kb) L1HS-L1PA3 elements elements gaining H3K4me3 in hNPCs and their respective IgG controls.

### Loss of SETDB1-mediated H3K9me3 results in transcriptional activation of L1s

To investigate the transcriptional consequences on L1s and other TEs after loss of H3K9me3 in hNPCs, we performed bulk RNA sequencing of SETDB1-CRISPRi and control hNPCs. We used an in-house 2 x 150 bp poly(A)-enriched stranded library preparation with a reduced fragmentation step to optimize read length for L1 analysis **(Fig. 4A)**. We obtained between 26 – 76 million uniquely mapped reads per sample. We used a stringent unique mapping approach to quantify expression of individual TE loci (21).

We found 3643 individual TE loci that were upregulated in SETDB1-CRISPRi hNPCs (DESeq2 LFC > 0, two-sided padj 0.05) confirming that many TEs are transcriptionally activated upon loss of H3K9me3 **(Fig. 4B)**. Strikingly, significantly upregulated TEs were enriched for L1s **(Fig 4C)**. When analysing L1-subfamilies we found that this enrichment of upregulated elements was limited to evolutionarily young L1s, including human-specific L1HS and hominoid-specific L1PA2-L1PA3s (**Fig. 4D-E)**. In total, we found 146 individual, full-length L1HS-L1PA3 elements being significantly upregulated **(Fig. 4D-E)**. We found a similar upregulation of L1s when using both SETDB1 gRNAs (**Sup fig. 5A-C**).

**Figure 5.**
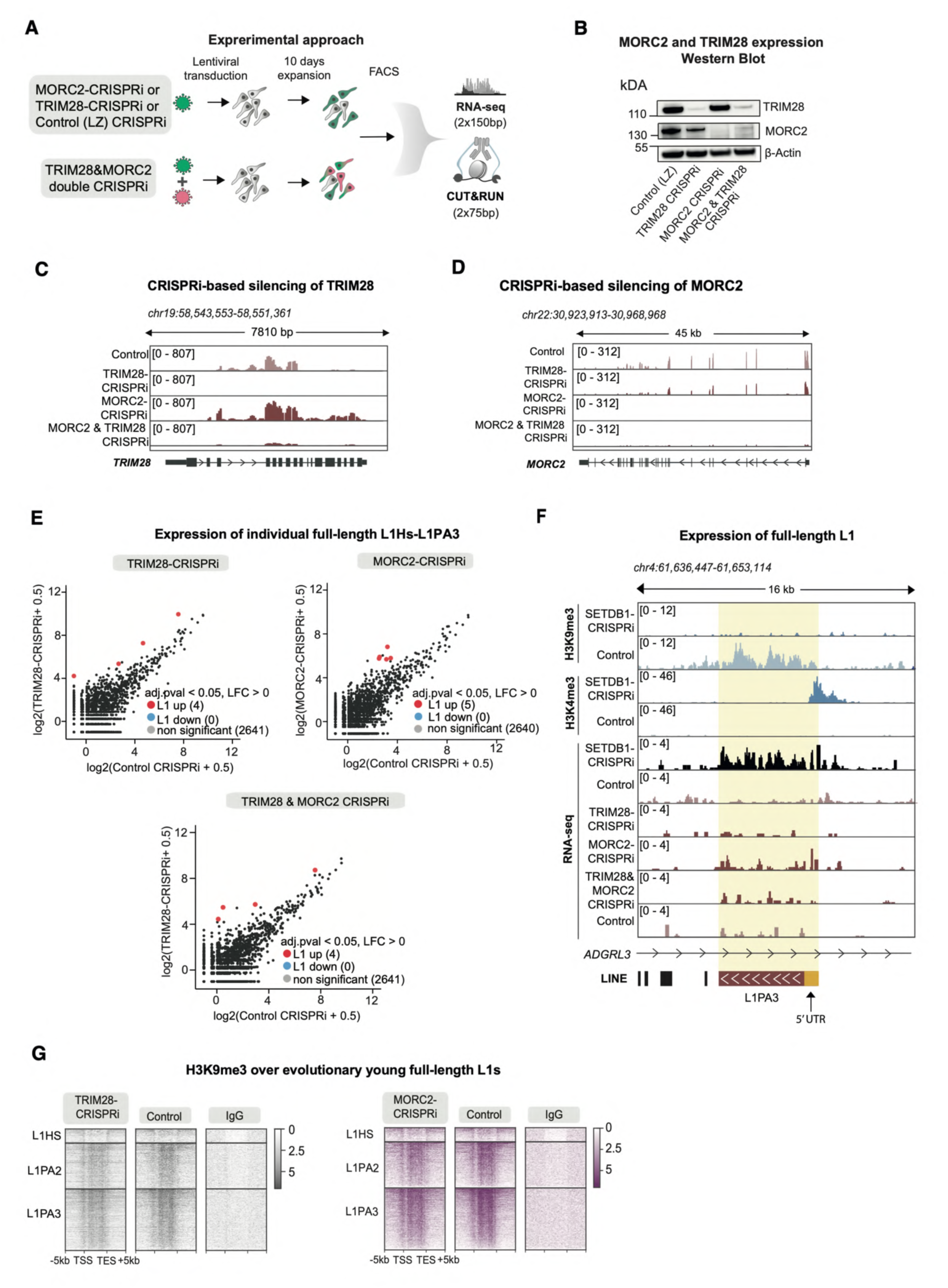
SETDB1 regulates evolutionary young, full-length L1 elements independently from MORC2 and TRIM28. **A** Workflow of H3K9me3 and H3K4me3 CUT&RUN and RNA sequencing analysis in TRIM28-CRISPRi, MORC2-CRISPRi, TRIM28&MORC2 double CRISPRi and control hNPCs. **B** Western blot analysis of TRIM28 and MORC2 protein levels in TRIM28-CRISPRi, MORC2-CRISPRi, TRIM28&MORC2 double CRISPRi and control hNPCs relative to ß-actin protein levels. **C** RPKM normalized genome browser tracks showing TRIM28 expression in TRIM28-CRIPSRi, MORC2-CRISPRi, TRIM28&MORC2 double CRISPRi and control hNPCs. **D** RPKM normalized genome browser tracks showing MORC2 expression in TRIM28-CRIPSRi, MORC2-CRISPRi, TRIM28&MORC2 double CRISPRi and control hNPCs **E** Mean plot of uniquely mapped bulk RNA sequencing reads for TRIM28-CRISPRi (n=3), MORC2-CRISPRi (n=3), TRIM28&MORC2 double CRISPRi (n=3) hNPCs compared to controls (n=3) over full-length (>6 kb) L1HS-L1PA3 elements. LFC > 0, two-sided padj < 0.05 calculated with DESeq2. **F** RPKM normalized genome browser tracks showing: CUT&RUN analysis in SETDB1-CRISPRi hNPCs of H3K9me3 (top) and H3K4me3 (middle). RNA expression of an L1PA3 element in SETDB1-CRISPRi, TRIM28-CRISPRi, MORC2-CRISPRi, TRIM28&MORC2 double CRISPRi and hNPCs compared to controls (bottom). **G** RPKM normalized heat maps showing CUT&RUN signal of H3K9me3 over full-length (> 6kb) L1HS to L1PA3 elements sorted by evolutionary age in MORC2-CRISPRi and TRIM28-CRISPRi hNPCs compared to their respective controls and non-targeting control IgG. Data from Pandiloski et al. (2024) (23) and Horvath et al. (2024) (28)

When comparing the number of reads transcribed in the same orientation as the L1 (in sense) to reads in the opposite direction (in antisense) we found that most of the transcription in these regions were in sense with regards to the L1s in SETDB1-CRISPRi hNPCs **(Fig 4F, Sup fig. 5D)**. This is consistent with the idea that the detected L1 transcripts come from the L1 promoter, rather than being a result of read-through transcription. We also investigated the genomic location of the L1s that were found to be upregulated in SETDB1-CRISPRi hNPCs. The majority of the upregulated L1s were found in intergenic regions of the genome **(Fig. 4G, Sup fig. 5E)**. Of those located within genes, the majority were found to be antisense to the host genes **(Fig. 4G, Sup fig. 5E)**. These findings also suggest that the upregulated L1 expression is a consequence of elevated L1 transcription, rather than being related to the expression of genes harbouring an L1 in their introns. In addition to L1s, we also found upregulation of other TEs, such as several ERV families and SVAs, which are know to be controlled by the KZNF-TRIM28 system that depends on SETDB1 **(Table S1.)** (28,50–52).

To further confirm that the the epigenetic status of L1 promoters is associated with an active state in SETDB1-CRISPRi hNPCs, we performed CUT&RUN analysis for the histone mark H3K4me3, which is associated with active promoters. Although this approach is limited to L1 elements where the H3K4me3 peak is broad enough to spread to the surrounding genomic context, we were able to detect 24 high-confidence H3K4me3 peaks located at the 5′ end of evolutionarily young, full-length L1s in SETDB1-CRISPRi NPCs (**Fig 4H-I, Sup fig. 5F**). Most of these loci were also transcriptionally activated and lost H3K9me3 (**Fig 4H-I, Sup fig. 5F**). Taken together, these results show that SETDB1-mediated H3K9me3 is essential to silence evolutionarily-young, full-length L1 elements in hNPCs.

### SETDB1 regulates the expression of evolutionarily young L1s independently from the HUSH/MORC2 corepressor and the KZNF-TRIM28 pathways

The observation that evolutionarily young L1s are activated by loss of H3K9me3 was surprising, as previous work has indicated that DNA methylation is the key repressive mechanism of L1s in neural cells (8,9,23,24). The two major pathways that control the transcription of TEs in association with SETDB1 and H3K9me3 are the KZNF-TRIM28 system and the human silencing hub (HUSH)/MORC2 co-repressor pathway (29,53–56). Previous studies in hNPCs have shown that loss of these pathways in somatic cells has only a limited effect on L1 expression (23,50). To confirm this, we performed lentiviral-based TRIM28-CRISPRi and MORC2-CRISPRi experiments in hNPCs, comparing the transcriptional activation of evolutionarily-young full-length L1s with that of SETDB1-CRISPRi hNPCs **(Fig. 5A)**. Ten days after lentiviral transduction, we verified the efficient silencing of MORC2 and TRIM28 using qRT-PCR, WB and RNA-seq **(Fig. 5B-D, Sup fig.6A)**. Silencing of TRIM28 or MORC2 resulted in upregulation of only 4 and 5 individual, full-length L1HS-L1PA3 elements, respectively **(Fig. 5E-F)**, which can be compared with 146 individual, full-length L1HS-L1PA3 elements being significantly upregulated after SETDB1 silencing (**Fig. 4D**). To verify that the lack of a transcriptional response of L1s upon silencing of TRIM28 or MORC2 was not due to a redundancy between these pathways, we performed a double CRISPRi-silencing experiment of these two genes (**Fig. 5A**). We verified the efficient double silencing of TRIM28 and MORC2 using qRT-PCR, WB and RNA-seq **(Fig. 5B-D, Sup fig.6A)**. Silencing of both MORC2 and TRIM28 did also not result in transcriptional activation of L1s, only 4 indivual full-length L1HS-L1PA3 elements were upregulated in this setting **(Fig. 5E-F)**. We also used our previously published H3K9me3 CUT&RUN data from TRIM28-CRISPRi and MORC2-CRISPRi hNPCs (23,28) to investigate if these genes are responsible for H3K9me3 at L1s in hNPCs. Gobal analysis following the inactivation of TRIM28 or MORC2 confirmed the expected loss of the H3K9me3 signal (Sup fig.6B-C). SETDB1-CRISPRi resulted in major loss of H3K9me3 peaks over full-length L1HS-L1PA3 elements **(Fig. 2I),** while MORC2-CRISPRi and TRIM28-CRISPRi had minor effects on H3K9me3 deposition over full-length L1HS-L1PA3 elements **(Fig. 5G).** Together these results show that in addition to the TEs that are repressed by HUSH and KZNF-TRIM28 and thus dependendent on SETDB1, SETDB1 also acts through alternative pathways to maintain L1 promoters with H3K9me3 leading to their transcriptional repression.

**Figure 6.**
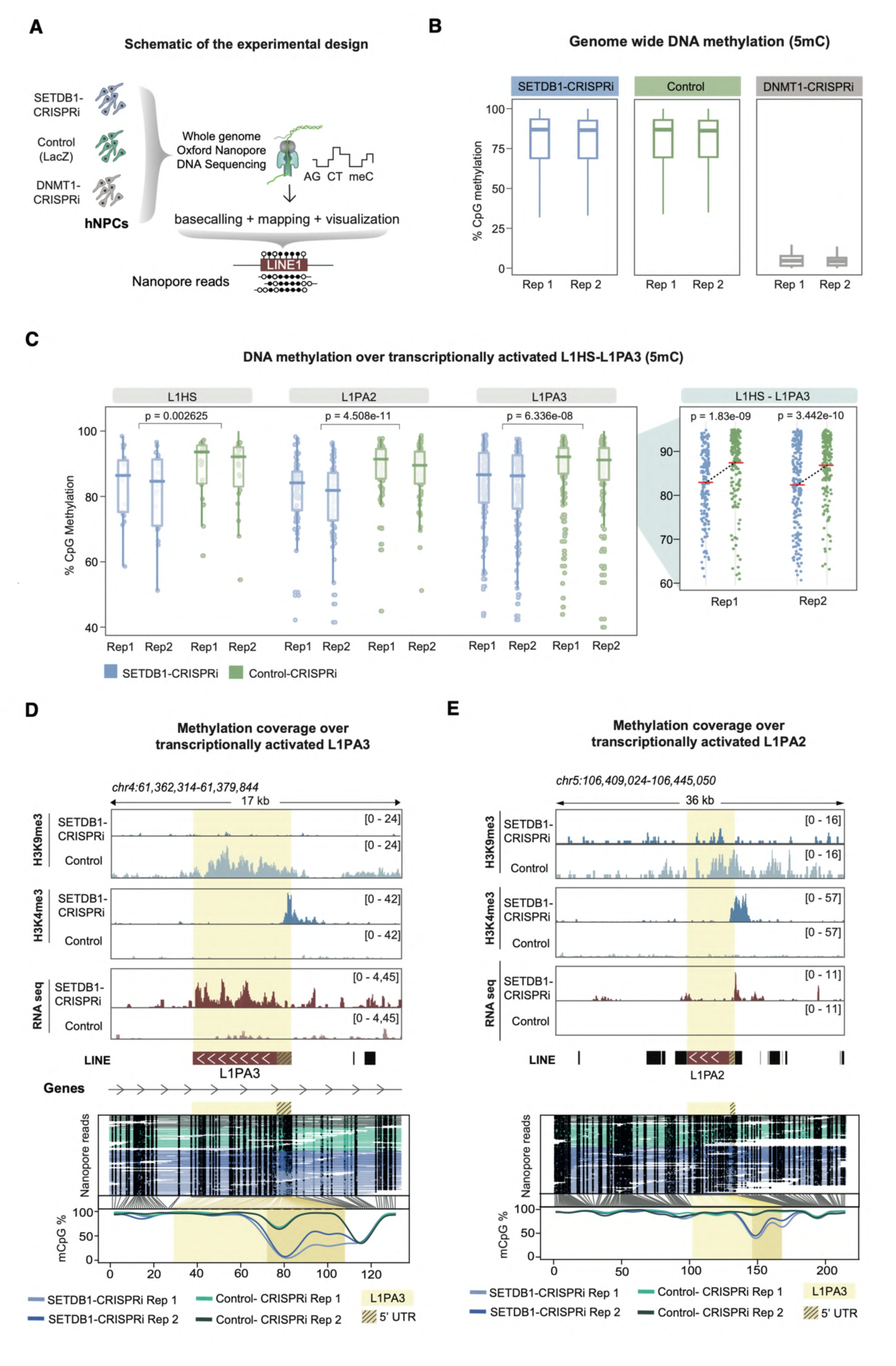
Loss of H3K9me3 results in loss of CpG DNA methylation at L1 promoters in hNPCs. **A** Schematics of whole genome ONT long-read DNA sequencing analysis in hNPCs **B** Box plots of the genome wide CpG DNA methylation status in SETDB1-CRISPRi (n=2), control (n=2) and DNMT1-CRISPRi (n=2) hNPCs. **C** Box plots of the methylation status over the promoter of transcriptionally activated L1HS-L1PA3 elements in SETDB1-CRISPRi (n=2) and control (n=2) hNPCs. Zoom-in panels indicating mean methylation levels (red line) per condition for L1HS-L1PA3 elements. The figure shows FL-L1s that were found to be upregulated in at least one of the two experimental conditions (two different gRNAs). This resulted in 241 FL-L1s being included in the analysis. n values: L1HS = 22; L1PA2 = 77; and L1PA3 = 142. **D&E** RPKM normalized genome browser tracks showing from top to bottom: CUT&RUN analysis of H3K9me3 and H3K4me3 in SETDB1-CRISPRi compared to control hNPCs, RNA expression of L1s in SETDB1-CRISPRi compared to controls hNPCs. ONT DNA reads across SETDB1-CRISPRi and control hNPCs. Black dots indicate methylated CpGs, and methylation coverage of the L1 element can be seen at the bottom. The L1 element is highlighted in yellow, while the 5’ UTR is marked with stripes.

### Loss of H3K9me3 results reduced CpG DNA methylation at L1 promoters

CpG DNA methylation of the L1 promoter has been described as the major mechanism for transcriptional silencing of L1 elements in somatic cells, including hNPCs (7–10,22,23). Therefore, we hypothesized that loss of H3K9me3 may have downstream effects on DNA methylation levels at L1s. To investigate the interplay between H3K9me3 and CpG DNA methylation, we performed genome-wide DNA methylation profiling using Oxford Nanopore Technologies (ONT) long-read DNA sequencing on SETDB1-CRISPRi and control hNPCs **(Fig. 6A)**. As an additional positive control, we used CRISPRi to silence DNMT1 (23,28), whose deletion in hNPCs results in a global loss of DNA methylation, including over L1s (8) (**Fig. 6A, Sup Fig 7A-B**).

**Figure 7.**
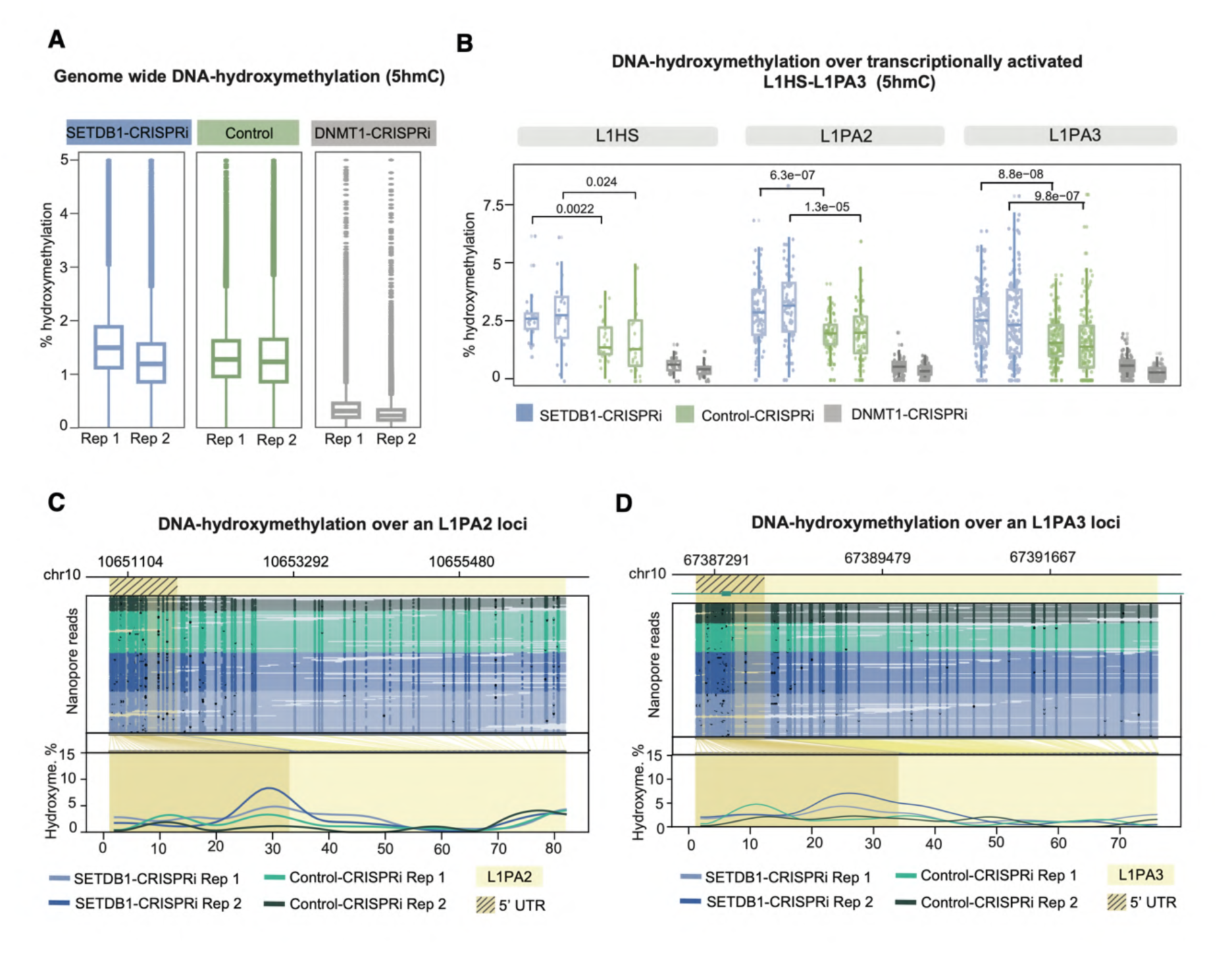
Loss of H3K9me3 results in gain of DNA hydroxymethylation. **A** Box plots of the genome wide DNA hydroxymethylation status in SETDB1-CRISPRi (n=2), control (n=2) and DNMT1-CRISPRi (n=2) hNPCs. **B** Box plots of the DNA hydroxymethylation status over the promoter of transcriptionally activated L1HS-L1PA3 elements in SETDB1-CRISPRi (n=2), control (n=2)and DNMT1-CRISPRi hNPCs. The figure shows FL-L1s that were found to be upregulated in at least one of the two experimental conditions (two different gRNAs, 241 FL-L1s being included in the analysis. N: L1HS = 22; L1PA2 = 77; and L1PA3 = 142). **C** Locus plot showing DNA hydroxymethylation over an upregulated L1PA2 in SETDB1-CRISPRi (blue, n=2) and control (green, n=2) hNPCs. Black dots indicate methylated CpGs, and methylation coverage of the L1 element can be seen at the bottom. The L1 element is highlighted in yellow, while the 5’ UTR is marked with stripes. **D** Locus plot showing DNA hydroxymethylation over an upregulated L1PA3 in SETDB1-CRISPRi (blue, n=2) and control (green, n=2) hNPCs. Black dots indicate methylated CpGs, and methylation coverage of the L1 element can be seen at the bottom. The L1 element is highlighted in yellow, while the 5’ UTR is marked with stripes.

We observed no major changes in global genome-wide DNA methylation between SETDB1-CRISPRi and control hNPCs, suggesting that DNA methylation patterns are generally unaffected by the loss of H3K9me3 over the 10-day timecourse of the experiment **(Fig. 6B)**. This contrasts with the silencing of DNMT1, which results in global DNA hypomethylation **(Fig. 6B)**. However, when we performed DNA methylation analysis of evolutionarily young L1s, we found that SETDB1-CRISPRi hNPCs exhibited lower levels of CpG DNA methylation compared to control hNPCs. In particular, L1s that were transcriptionally activated in hNPCs showed significantly lower levels of DNA methylation over their 5’ promoter region **(Fig. 6C, Sup Fig.7C-E)**. For example, we found full-length human- and hominoid-specific L1 elements that were transcriptionally activated in SETDB1-CRISPRi hNPCs, as determined by bulk RNA sequencing and CUT&RUN profiling of the active promoter mark H3K4me3, and also lost DNA methylation **(Fig. 6D-E)**. When investigating DNA methylation over all full-length L1s and those which did not get transcriptionally activated we found that some L1s in these groups also loose DNA methylation, highlighting the challanges to detect the RNA signal for some of the evolutionary young L1s (Sup Fig.7F-I). Taken together, these results demonstrate that the loss of SETDB1-mediated H3K9me3 in human somatic cells leads to subsequent loss of DNA methylation and gain of H3K4me3 methylation, resulting in transcriptional activation of L1s.

The loss of DNA methylation could result from an active process involving ten-eleven translocation (TET) enzymes, which is characterized by the conversion of methylcytosine to hydroxymethylcytosine (57). Alternatively, it may result from a passive process associated with an inability to attract DNMT1, which is characterized by the loss of methylcytosine. Therefore, we investigated the presence of DNA hydroxymethylation in SETDB1-CRISPRi and control hNPCs, as determined from the ONT data, using DNMT1-CRISPRi as a negative control. While we did not see global changes in DNA hydroxymethylation at a genome-wide level in SETDB1-CRISPRi compared to control cells, our analysis revealed that transcriptionally activated full-length L1s gained DNA hydroxymethylation over their promoter (Fig. 7B-D, Sup Fig.8A-F). This suggests that the loss of DNA methylation at L1s with SETDB1-CRISPRi occurs via an active process involving TET enzymes. In contrast, DNMT1-CRISPRi NPCs do not exhibit an increase in DNA hydroxymethylation at these elements (Fig. 7B). This is consistent with passive loss of DNA methylation resulting from the loss of DNMT1 activity. Thus, these results demonstrate that the loss of DNA methylation specifically over the promoter region of L1s in SETDB1-CRISPRi NPCs is an active process likely mediated by the recruitment of TET enzymes.

## Discussion

Here we study the consequences of loss of H3K9me3 heterochromatin in hNPCs, a cell type that allows us to model some aspects of heterochromatin maintenance in a human setting, including the presence of H3K9me3 and DNA methylation at L1s (8,23). Upon CRISPRi-based silencing of the H3K9me3 methyltransferase SETDB1 in hNPCs, we found a major loss of H3K9me3 peaks, including those over L1s and other TEs. Moreover, we observed a general reorganization of H3K9me3 domains, including the appearance of new ectopically large heterochromatin domains, as detected by both CUT&RUN analysis and confocal microscopy. While we do not fully understand the mechanism behind this phenomenon, previous studies have reported that a global loss of H3K9me3 leads to rearrangements of the epigenome and the gain of H3K9me3 at specific loci mediated by other H3K9me3 methyltransferases such as SUV39H1/2 (58–60). However, despite this global heterochromatin reorganization, SETDB1-CRISPRi hNPCs remained viable, proliferative, and express appropriate levels of NPC marker genes, suggesting that the presence of H3K9me3 at TEs is not critical for maintaining somatic cell identity and viability, at least under the experimental conditions in this study.

L1s are the only autonomously mobilizing family of TEs in humans (61) and can impact genome stability via several mechanisms, including new somatic retrotransposition events and alterations of transcriptional networks where transcriptionally active L1s may impact the genome both in *cis* and *trans* (21,62–66). L1s are therefore mostly transcriptionally silenced in somatic human cells. Our data demonstrate that H3K9me3 and SETDB1 play a critical role in this process. In the absence of SETDB1 and H3K9me3, DNA methylation is lost over the promoter of evolutionary young L1s, which correlates with their transcriptional activation. This is in line with a model where the deposition of H3K9me3 at the L1 promoter is needed to maintain DNA methylation (67–69). The best understood mechanisms for deposition of H3K9me3 on TEs are the KRAB zinc finger proteins (KZNF)-TRIM28 system and the HUSH pathway. However, the control of evolutionary young L1s by KZFNs remains poorly understood. A previous study found that ancestral L1PA3 elements underwent structural sequence changes that allowed them, and subsequent younger L1 families, to escape silencing by ZNF93, which otherwise targets L1s (70). Although other KZNFs have been reported to bind to young L1s (31), these have not been studied in detail and appear not to be active in NPCs, as evidenced by the results of our TRIM28-CRISPRi experiments. The HUSH pathway can target evolutionary young L1s (56,71), but recent data from our labs show that this is restricted to transcriptionally active elements that are not covered by DNA methylation at steady-state, as is the case for almost all L1s in hNPCs (8,9,23). Thus, the pathway by which H3K9me3 is maintained over evolutionarily-young L1s resulting in their silencing in somatic cells remains to be fully elucidated. However, our data show that the deposition of H3K9me3 at L1s is necessary to maintain DNA methylation over these elements in somatic cells. This likely occurs by counteracting the demethylation activity of TET enzymes. Thus, our data demonstrate that DNMT1 alone is insufficient for maintaining methylation patterns at the promoter of evolutionary young L1s.

The loss of H3K9me3 and heterochromatin is a hallmark of aging (72). Experiments conducted across diverse organisms and tissues have provided evidence of age-related heterochromatin loss and there is a gradual loss of H3K9me3 during the aging process in human model systems (73–76). While our H3K9me3 deficient hNPCs have obvious limitations as a model for human brain ageing, our results show that a key event following the loss of H3K9me3 maintenance is the transcriptional activation of L1s. In line with this, a number of recent studies have shown that increased L1 expression correlates with senescence and other age-related cellular phenotypes, and suggest that aberrant transcription of L1s may be associated with ageing and related disease processes (77–81).

The transcriptional activation of L1s is likely to have consequences, as evolutionarily young, intact L1 elements have the potential to retrotranspose and thus can cause deleterious mutations and structural variants (11,82–85). In addition, several recent studies have shown that L1 transcription itself can affect multiple cellular pathways both in *cis* and in *trans*. For example, we have recently shown that when DNA methylation is experimentally removed in human NPCs, transcriptionally activated L1 elements act as alternative promoters, affecting the expression levels of many protein-coding genes (8,21). These observations suggest that the aberrant activation of L1s could lead to widespread dysregulation of gene regulatory networks. In line with this, we observed in the current study that genes located in proximity of full-length L1HS-L1PA3 are upregulated upon SETDB1-CRISPRi **(Sup Fig. 9A)**. This supports the idea that L1 elements regulated by H3K9me3 in hNPCs can have a regulatory effect on nearby gene expression. Furthermore, L1 transcripts and L1-derived peptides can be sensed by the cellular machinery to trigger an inflammatory response. For example, L1 expression has been reported to be increased in senescent cells and to activate an interferon response (78,79,86,87). Thus, transcriptional activation by L1s in aged cells, as a consequence of H3K9me3 loss, may contribute to the ageing process via multiple mechanisms. However, we did not observe an interferon response in SETDB1-CRISPRi hNPCs, which may be explained by the cell type used in this study **(Sup Fig. 9B)**.

In conclusion, our results provide a unique insight into the consequences of loss of H3K9me3 in human somatic cells. SETDB1-mediated H3K9me3 is not required to maintain cellular identity and viability, but rather provides a critical mechanism to repress evolutionarily young L1 elements.

## Data availability

All data needed to evaluate the conclusions in the paper are present in the paper and/or the Supplementary Materials. The sequencing data presented in this study has been deposited at the GEO superseries GSE292899. Reviewer access to the data: GSE294699 (https://www.ncbi.nlm.nih.gov/geo/query/acc.cgi?acc=GSE294699; token: stgpoewcllshdkz) and GSE292899 (https://www.ncbi.nlm.nih.gov/geo/query/acc.cgi?acc=GSE292899; token: srwfscggjlepdiz). Original code has been deposited at GitHub and is publicly available at git@github.com:Molecular-Neurogenetics/SETDB1_Karlsson_2025.git.

## Acknowledgements

We would like to thank S. Henikoff for providing reagents and U. Jarl, for technical assistance. We acknowledge Clinical Genomics Lund, SciLifeLab, and Center for Translational Genomics (CTG) Lund University for providing expertise and service with sequencing and analysis. We are grateful to all members of the Jakobsson laboratory.

## Author contributions

Design and interpretation: All authors. Conceptualization: O.K., N.P., and J.J. Experimental research: O.K., A.A., V.H., P.J. and J.G.J. Bioinformatics: N.P. and R.G. Writing—original draft: O.K., N.P. and J.J. Writing—review and editing: All authors.

## Supplementary Figures

**Supplementary Figure 1.**
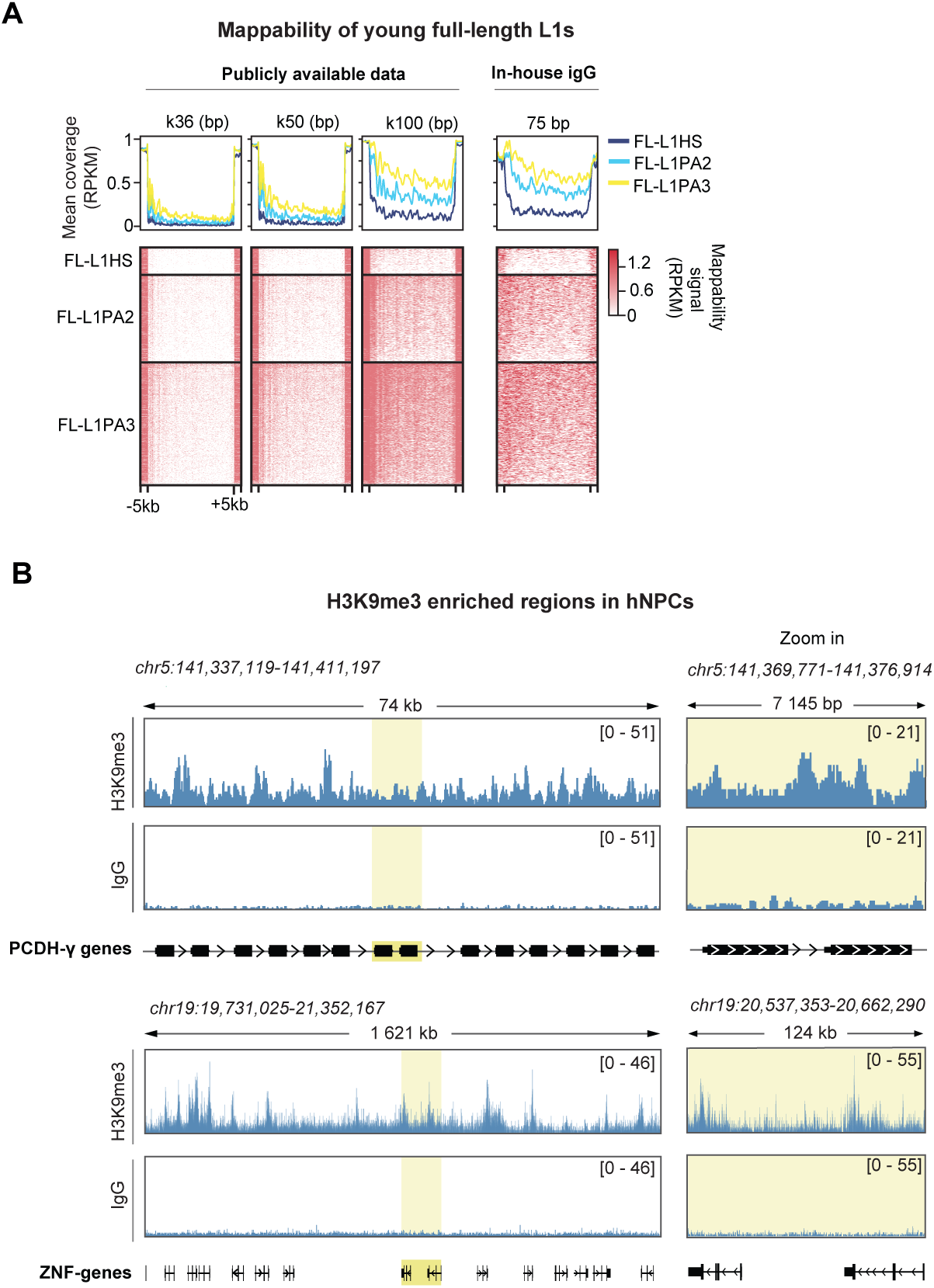
H3K9me3 in hNPCs localizes to regions with well-established H3K9me3-enrichment. **A** Mappability analysis of young full-length L1HS to L1PA3 using 36bp, 50bp and 100bp uniquely mappable kmers obtained from publicly available dataset (10.1093/nar/gky677), and an example of an in-house IgG experiment. **B** CUT&RUN analysis of H3K9me3 in hNPCs over genomic regions with well-established H3K9me3-enrichment; PCDH-ɣ cluster (top); KRAB-ZNF cluster (bottom).

**Supplementary Figure 2.**
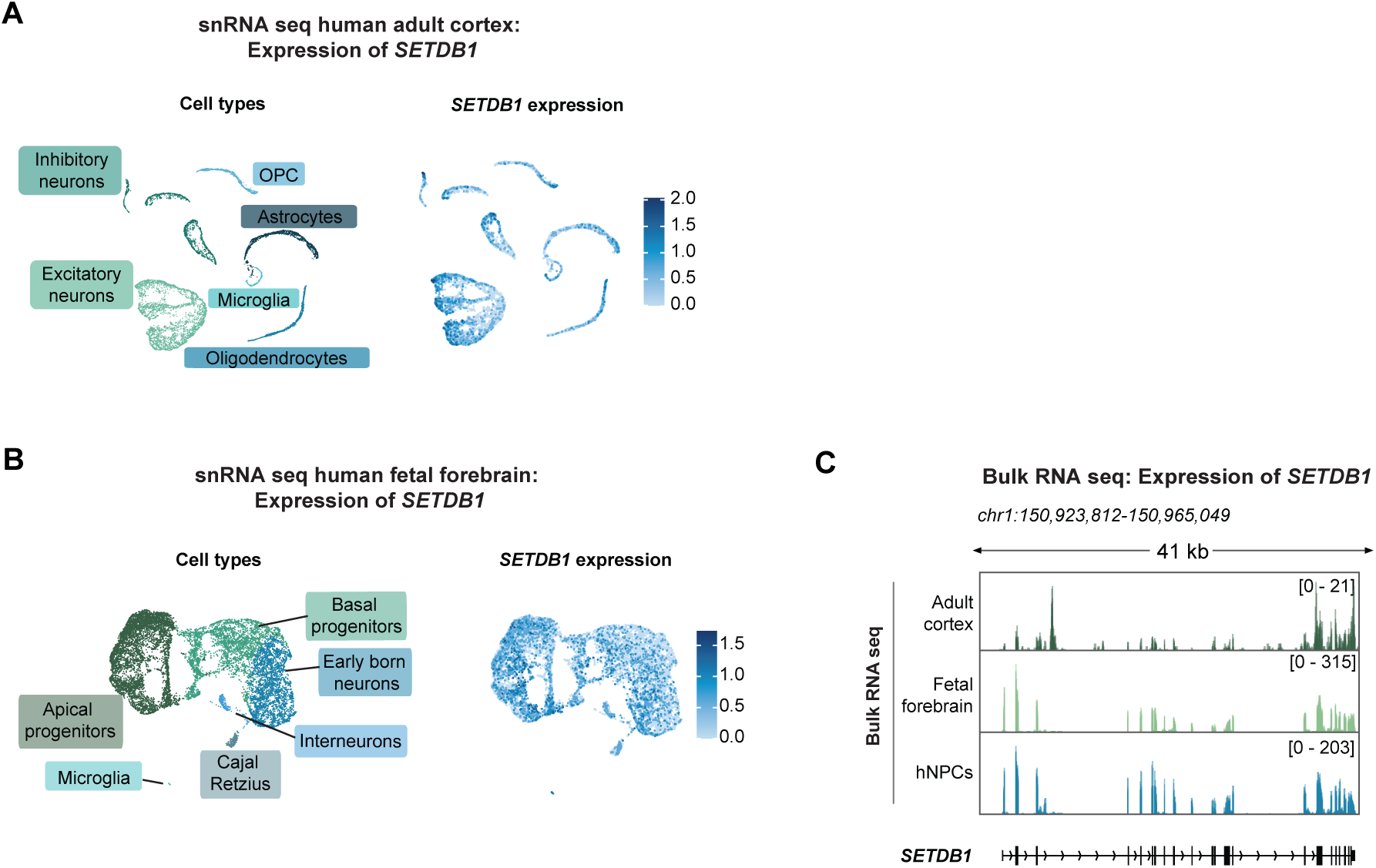
SETDB1 is expressed in the adult and developing human brain and hNPCs. **A-B** UMAP of snRNA-seq data showing SETDB1 expression in (A) human adult cortical tissue and (B) human fetal forebrain tissue. **C** Genome browser tracks showing bulk RNA sequencing data of SETDB1 expression in the human adult cortex, human fetal forebrain, and hNPCs. Data from Garza *et al.* (2023) (20).

**Supplementary Figure 3.**
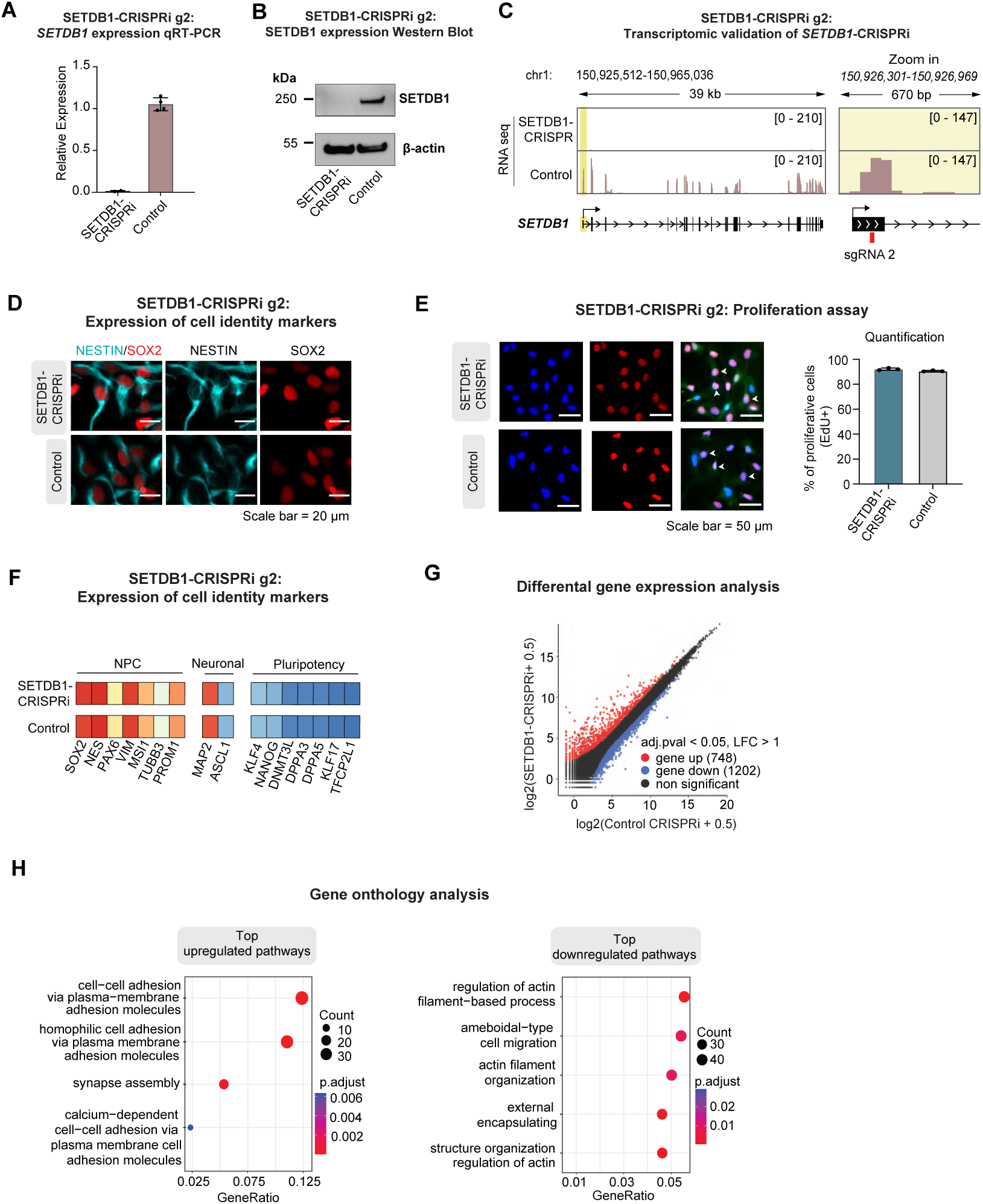
Validation of SETDB1-CRISPRi using an alternative guide RNA. **A** qRT-PCR analysis of SETDB1 expression after CRISPRi g2-based silencing in hNPCs. Results are shown as means with standard deviation (*n*=2 and *n*=4 respectively). **B** Western blot analysis of SETDB1 protein levels relative to β-actin protein levels in SETDB1-CRISPRi g2 and control hNPCs. **C** RPKM normalized genome browser tracks showing expression of *SETDB1* in SETDB1-CRISPRi g2 and control hNPCs as determined by bulk RNA sequencing **D** Immunocytochemistry of NESTIN (cyan) and SOX2 (red) in SETDB1-CRISPRi g2 and control hNPCs. Scalebar = 20 μm. **E** EdU proliferation assay showing cell proliferation in SETDB1-CRISPRi g2 and Control hNPCs. Scale bar=50 μm. **F** Heat map of NPC, neuronal and pluripotency gene marker mean expression in SETDB1-CRISPRi g2 (*n*=2) and control (*n*=4) hNPCs as determined by bulk RNA sequencing. **G** Mean plot showing differential gene expression in SETDB1-CRISPRi (*n*=4) compared to control (*n*=4) hNPCs. LFC >1, padj < 0.05 calculated with DESeq2. **H** Gene-ontology overrepresentation test of significantly differentially expressed genes (|LFC| > 1) associated with the selected top upregulated terms (left) and the selected top downregulated terms (right) in SETDB1-CRISPRi compared to control hNPCs.

**Supplementary Figure 4.**
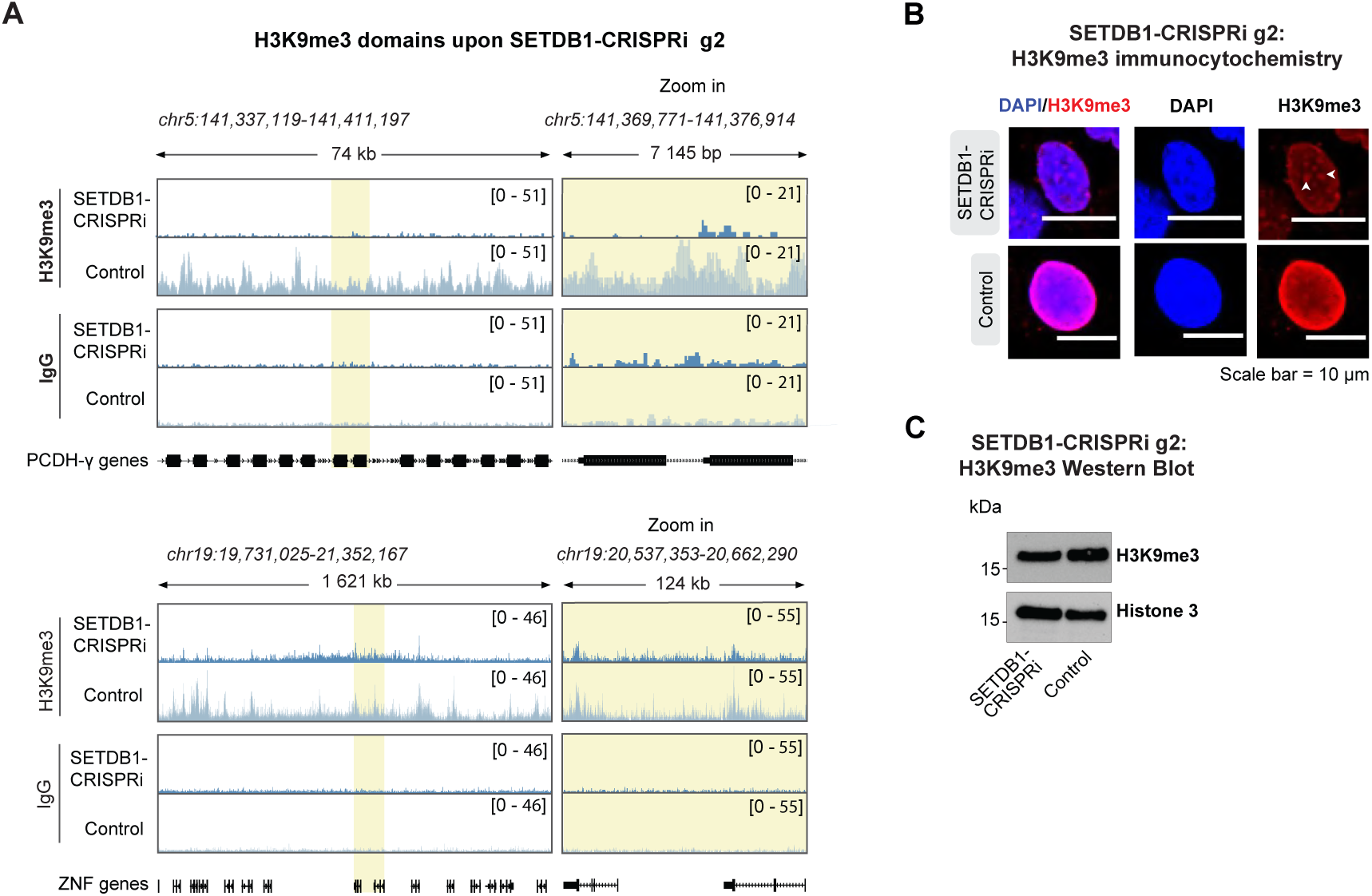
H3K9me3 enrichment over genomic regions. **A** CUT&RUN analysis of H3K9me3 over genomic regions with well-established H3K9me3-enrichment in SEDTB1-CRISPRi and control hNPCs **B** Immunocytochemistry of H3K9me3 (red) and nuclear marker DAPI (blue) in SETDB1-CRISPRi g2 and control hNPCs. H3K9me3 foci are indicated with white arrows. Scalebar = 10 μm. **C** Western blot analysis of H3K9me3 levels relative to Histone 3 in SETDB1-CRISPRi g2 hNPCs compared to control.

**Supplementary Figure 5.**
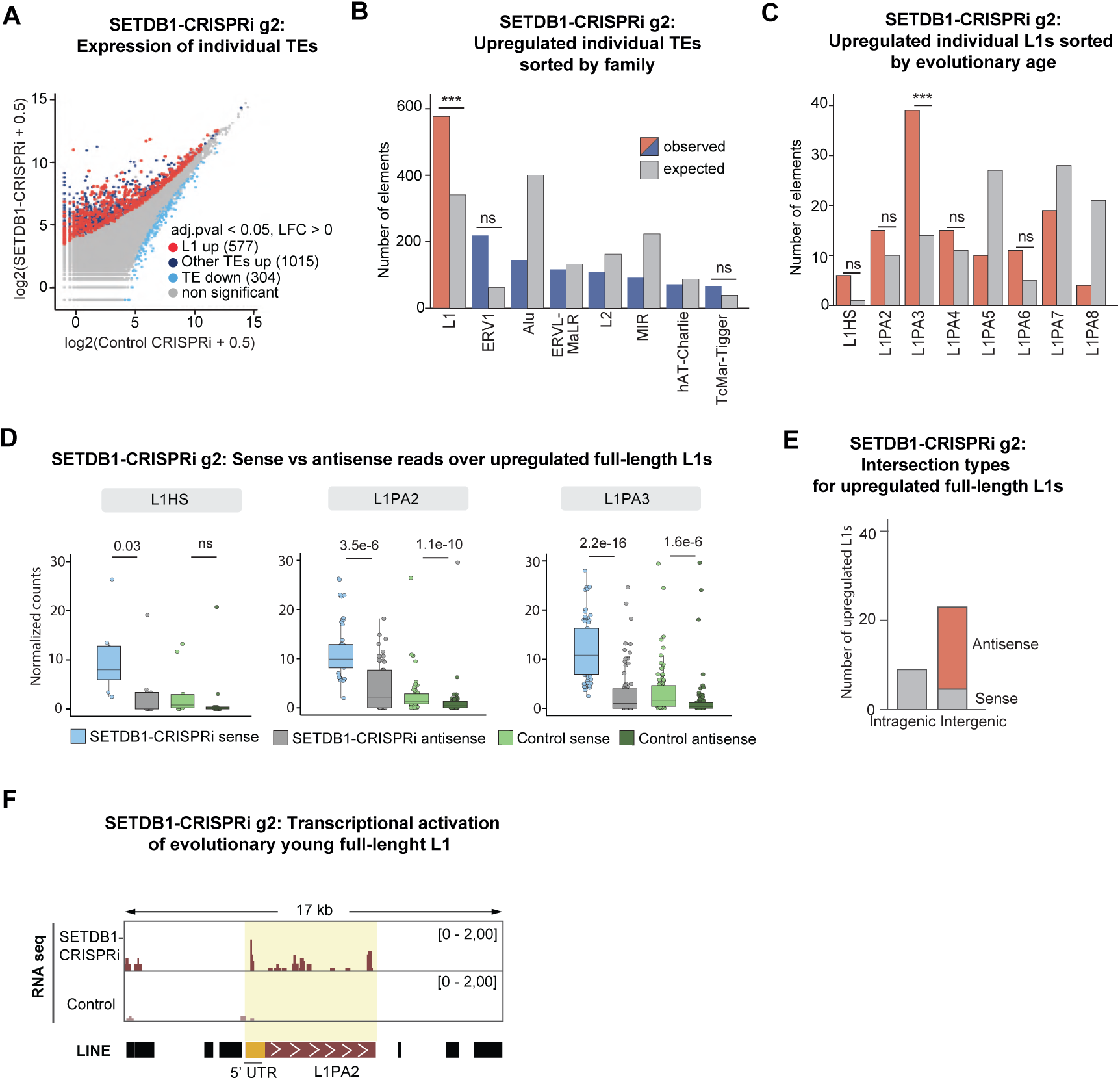
Validation of transcriptional activation of L1s upon SETDB1-CRISPRi using an alternative guide RNA. **A** Mean plot of uniquely mapped bulk RNA sequencing reads for SETDB1-CRISPRi g2 hNPCs (*n*=2) compared to control (*n*=2) over individual TEs categorized into L1 elements and other TEs. Log_2_ fold change (LFC) >0, padj 0.05 calculated with DESeq2. **B** Observed vs expected number of individual transcriptionally activated elements per TE family in SETDB1-CRISPRi g2 hNPCs (n= 2) compared to control (n= 2). P-values were calculated using bootstrap-based empirical two-sided p-value calculation. **C** Observed vs expected number of transcriptionally activated L1 elements sorted by evolutionary age in SETDB1-CRISPRi g2 *(n*=2) compared to control hNPCs *(n*=2). Pvalues were calculated using bootstrap-based empirical pvalue calculation. **D** Normalized read counts in sense and antisense for upregulated, full-length L1HS-L1PA3 elements in SETDB1-CRISPRi g2 (*n*=2) vs control hNPCs (*n*=4). Statistical significance calculated with Mann-Whitney U test. **E** Number, genomic location and direction of upregulated full-length (>6 kb) L1HS-L1PA3 elements in SETDB1-CRISPRi g2 (n=2) compared to control hNPC (n=2) ***F*** RPKM normalized genome browser tracks showing RNA-seq in SETDB1-CRISPRi g2 and control hNPCs.

**Supplementary Figure 6.**
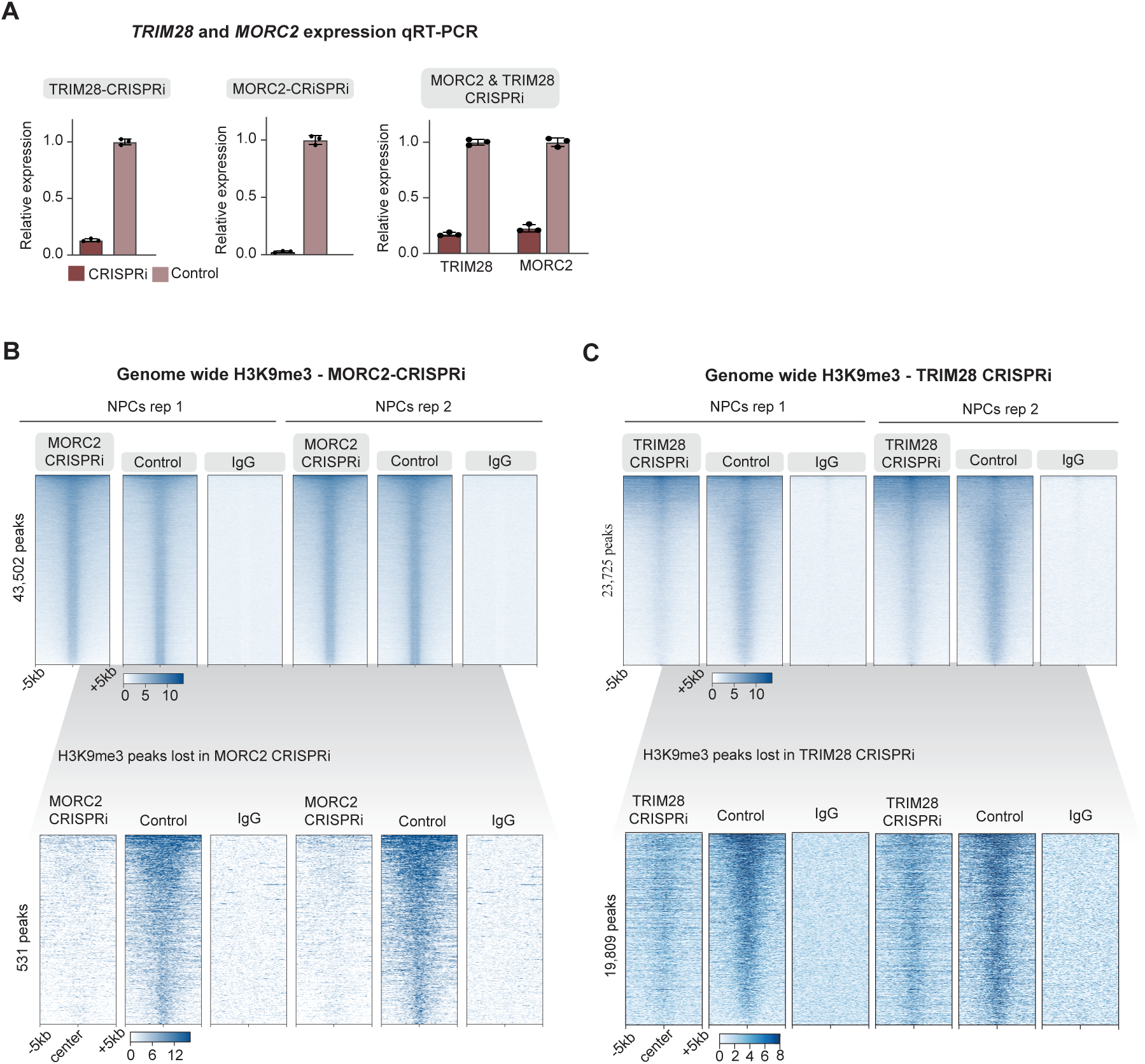
SETDB1 regulates evolutionary young, full-length L1 elements independently from MORC2 and TRIM28. **A** qRT-PCR analysis of TRIM28 and MORC2 expression after CRISPRi based silencing in hNPCs. Results are shown as means with standard deviation (*n*=3). **B** RPKM normalized heat maps showing H3K9me3 CUT&RUN signal in MORC2-CRISPRi hNPCs genome-wide (top) and over genomic regions where H3K9me3 peaks were lost (bottom) compared to their respective controls and non-targeting control IgG. Data from Pandiloski et al. (2024) (23)**. C** RPKM normalized heat maps showing H3K9me3 CUT&RUN signal in TRIM28-CRISPRi hNPCs genome-wide (top) and over genomic regions where H3K9me3 peaks were lost (bottom) compared to their respective controls and non-targeting control IgG. Data from Horvath et al. (2024) (26).

**Supplementary Figure 7.**
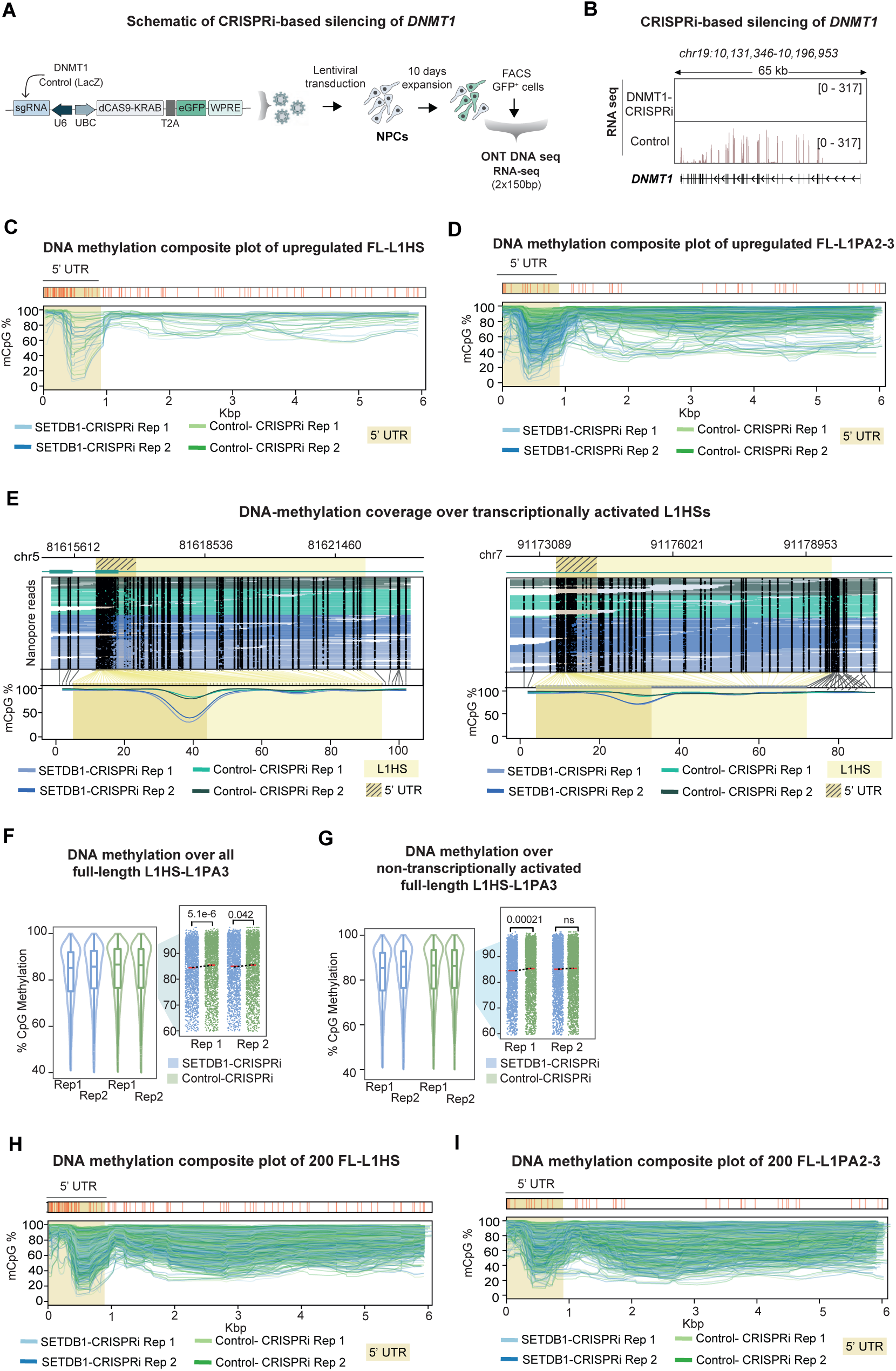
DNA methylation analysis over evolutionary young L1s. **A** Workflow for CRISPRi-based silencing of *DNMT1* in hNPCs and downstream analysis. **B** RPKM normalized genome browser tracks showing *DNMT1* expression in DNMT1-CRISPRi and control hNPCs. **C** Composite DNA methylation profiles of upregulated full-length L1HS in SETDB1-CRISPRi (blue, n=2) and control (green, n=2) hNPCs. The 5’ UTR is highlighted in yellow. **D** Composite DNA methylation profiles of upregulated full-length L1PA2 and L1PA3 in SETDB1-CRISPRi (blue, n=2) and control (green, n=2) hNPCs. The 5’ UTR is highlighted in yellow. **E** Locus plot showing DNA methylation over upregulated L1HS in SETDB1-CRISPRi (blue, n=2) and control (green, n=2) hNPCs. Black dots indicate methylated CpGs, and methylation coverage of the L1 element can be seen at the bottom. The L1 element is highlighted in yellow, while the 5’ UTR is marked with stripes. **F** Box plots of the methylation status over the promoter of all full-length L1HS, L1PA2 and L1PA3 elements in SETDB1-CRISPRi (n=2) and control (n=2) hNPCs. Zoom-in panels indicating mean methylation levels (red line) per condition for L1HS-L1PA3 elements. n values: L1HS = 306; L1PA2 = 972; L1PA3 = 1367. **G** Box plots of the methylation status over the promoter of full-length non-transcriptionally activated L1HS, L1PA2 and L1PA3 elements in SETDB1-CRISPRi (n=2) and control (n=2) hNPCs. Zoom-in panels indicating mean methylation levels (red line) per condition for L1HS-L1PA3 elements. n values: L1HS = 284; L1PA2 = 895; L1PA3 = 1225. **H** Composite DNA methylation profiles of 200 randomly picked full-length L1HS elements in SETDB1-CRISPRi (blue, n=2) and control (green, n=2) hNPCs. The 5’ UTR is highlighted in yellow. **I** Composite DNA methylation profiles of 200 randomly picked fulllength L1PA2-L1PA3 elements in SETDB1-CRISPRi (blue, n=2) and control (green, n=2) hNPCs. The 5’ UTR is highlighted in yellow.

**Supplementary Figure 8.**
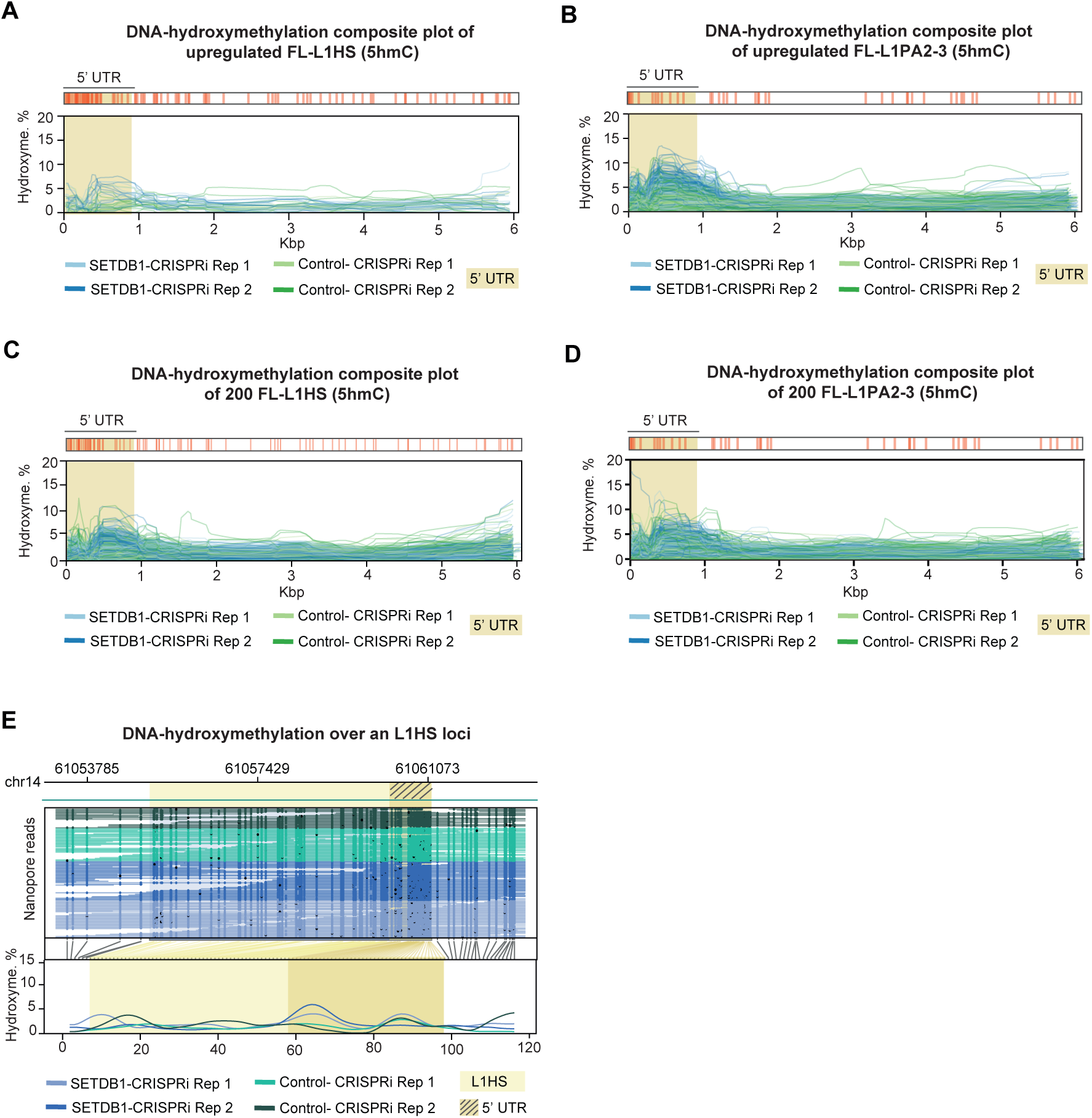
DNA-hydroxymethylation over evolutionary young L1s. **A** Composite DNA hydroxymethylation profiles of upregulated full-length L1HS in SETDB1-CRISPRi (blue, n=2) and control (green, n=2) hNPCs. The 5’ UTR is highlighted in yellow. **B** Composite DNA hydroxymethylation profiles of upregulated full-length L1PA2 and L1PA3 in SETDB1-CRISPRi (blue, n=2) and control (green, n=2) hNPCs. The 5’ UTR is highlighted in yellow. **C** Composite DNA hydroxymethylation profiles of 200 randomly picked full-length L1HS elements in SETDB1-CRISPRi (blue, n=2) and control (green, n=2) hNPCs. The 5’ UTR is highlighted in yellow. **D** Composite DNA hydroxymethylation profiles of 200 randomly picked full-length L1PA2-L1PA3 elements in SETDB1-CRISPRi (blue, n=2) and control (green, n=2) hNPCs. The 5’ UTR is highlighted in yellow. **E** Locus plot showing DNA hydroxymethylation over upregulated L1HS in SETDB1-CRISPRi (blue, n=2) and control (green, n=2) hNPCs. Black dots indicate methylated CpGs, and methylation coverage of the L1 element can be seen at the bottom. The L1 element is highlighted in yellow, while the 5’ UTR is marked with stripes.

**Supplementary Figure 9.**
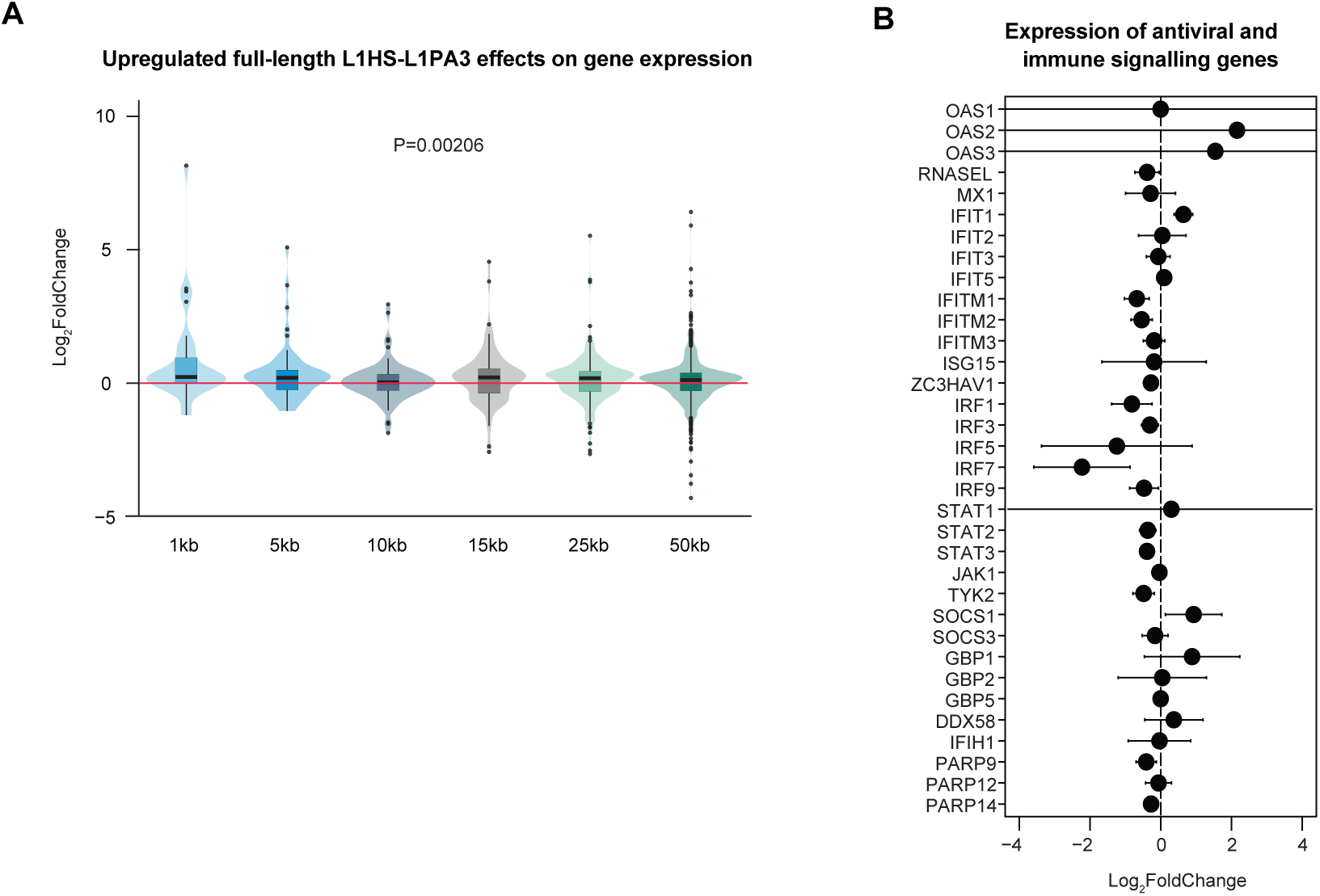
The effect of upregulated evolutionary young, full-length L1s on nearby gene expression. **A** Violin plots showing the effect of upregulated full-length L1HS-L1PA3 elements on nearby gene expression (2-50kb) upon SETDB1-CRISPRi in hNPCs. pvalue calculated with oneway ANOVA. **B** Differential expression analysis of selected antiviral and immune signaling genes in SETDB1-CRISPRi hNPCs (*n*=4) compared to control (*n*=4). LFC values are shown ± LFC standard error calculated by DESeq2.

**Supplementary Table 1.** Top 20 upregulated individual TEs upon SETDB1-CRISPRi. Table showing top 20 differentially expressed individual TEs as determined by bulk RNA sequencing in SETDB1-CRISPRi (*n*=4) and control (*n*=4) hNPCs. LFC and padj calculated with DESeq2.

| Individual TE | Log2FoldChange | p-adjusted value |
| --- | --- | --- |
| MER39B_dup439 | 12.5660730750503 | 1.8427529832473e-26 |
| L2c_dup54301 | 12.2209237438216 | 4.09869177917656e-25 |
| MER34B_dup283 | 11.7388139351731 | 2.93429232239475e-24 |
| MLT1B_dup6701 | 11.1359864402149 | 4.05191990872606e-19 |
| MIRb_dup150062 | 10.3913791660113 | 9.6211096384187e-19 |
| MIRb_dup83332 | 10.3742843350223 | 4.39773387345283e-17 |
| AluSx1_dup13234 | 10.3463773877304 | 2.18345277280823e-20 |
| L2c_dup54302 | 10.0803146120823 | 2.89355744423237e-16 |
| SVA_D_dup667 | 10.0527688112821 | 8.80024954039596e-19 |
| MIR_dup66266 | 9.91282533692722 | 1.32675089321779e-16 |
| Tigger1_dup11477 | 9.86311883738296 | 2.10590058343799e-18 |
| LTR10C_dup335 | 9.64204668384122 | 1.93395417652645e-17 |
| SVA_E_dup244 | 9.57541896859938 | 2.85991551525974e-12 |
| LTR12C_dup2419 | 9.34014514735954 | 8.07950807168578e-14 |
| THE1C_dup7048 | 9.17288181094644 | 1.27595527171481e-15 |
| LTR10C_dup29 | 9.03749224093489 | 6.18977818132119e-15 |
| L2c_dup96897 | 8.97118041487722 | 9.18595524144081e-15 |
| THE1B_dup16084 | 8.73842550859459 | 5.00823323103967e-14 |
| LTR79_dup800 | 8.51355207135647 | 7.37068570999639e-13 |
| THE1C_dup7049 | 8.49481998710542 | 5.63699471444779e-13 |

